# Assembly factors chaperone rRNA folding by isolating helical junctions that are prone to misfolding

**DOI:** 10.1101/2020.09.29.318998

**Authors:** Haina Huang, Katrin Karbstein

## Abstract

While RNAs are known to misfold, the underlying molecular causes remain unclear, and focused on alternative secondary structures. Conversely, how RNA chaperones function in a biological context to promote folding beyond duplex annealing, remains unknown. Here we show in a combination of DMS-MaPseq, structural analyses, biochemical experiments, and yeast genetics that three-way junctions are prone to misfolding during assembly of the small ribosomal subunit *in vivo*. We identify ubiquitous roles for ribosome assembly factors in chaperoning their folding by preventing the formation of tertiary interactions. In the absence of these assembly factors, tertiary interactions kinetically trap misfolded three-way junctions, thereby blocking further progress in the assembly cascade. While these protein chaperones act indirectly by binding the interaction partners, our analyses also suggest direct roles for snoRNAs in binding and chaperoning three-way junctions during transcription. This work furthermore shows that the dissociation of assembly factors renders reversible folding steps irreversible, thereby setting up a timer that regulates not just the flux of assembly but also dictates the propensity of misfolded intermediates to escape quality control. Finally, the data demonstrate how RNA chaperones act locally to unfold specific tertiary interactions, in contrast to protein chaperones, which globally unfold misfolded proteins.

## Introduction

Ribosomal RNA misfolds *in vitro (1-4)*, but folds correctly *in vivo*, where ∼ 200 factors promote its assembly(5-7), indicating a role for these factors in chaperoning RNA folding that remains largely uncharted. While the propensity of RNA to misfold is well known, even for small RNAs, its causes remain underexplored, especially for large RNAs such as the ribosome. Thus, it is also unclear how assembly factors could promote native structure; while a lot of attention has been paid to the role of alternative secondary structures in causing misfolding of RNA(8-17), less is known about misfolding of junctions or tertiary structure elements(10). This is likely because they are harder to probe, especially in the context of large RNAs such as rRNAs, and their perturbation is not as easily programmable as for secondary structures. Nonetheless, the observation that tertiary structures can stabilize both natively folded and misfolded structures(18), suggests the importance of specifying tertiary contacts for native or non-native folding. Finally, whether structure is formed (and misformed) at the local level, via individually misfolded elements, or rather globally, via entirely alternative sets of interactions remains unknown.

Here we address these questions using a tertiary interaction in the small ribosomal subunit head as a model system. The head is particularly prone to misfolding *(1-4)*, perhaps not surprising, given its unique challenge of needing to adopt multiple structures during translation(19-22). This tertiary interaction is formed by docking of a helical loop (l31) into a 3-way helical junction (j34) to form the subunit’s P-site (**Figure 1A-B)**. Using a combination of DMS-MaPseq, genetics, structural and biochemical analyses, we demonstrate a propensity for the 3-way junction to misfold and show that the misfolded intermediate becomes kinetically trapped once the tertiary interaction with l31 is formed. Moreover, we show how a succession of snoRNAs and assembly factors (AFs) binds l31, thereby preventing its contact with the 3-way junction. In the absence of this tertiary interaction, the 3-way junction can re-fold to the native state. Thus, these AFs chaperone folding of the 3-way junction indirectly by binding its interaction partner and thereby isolating the 3-way junction. Using these insights, we reanalyze previous *in vitro* folding data and the structures of assembly intermediates from cells. These analyses indicate that 3-way junctions are common bottlenecks for folding of ribosomal RNA, and also suggest general roles for snoRNAs and AFs in chaperoning the folding of these junctions by isolating them from interacting partners. Thus, these findings suggest an unappreciated role for 3-way junctions in the folding of complex structured RNAs, and a general mechanistic basis for the function of RNA chaperones. Lastly, our data indicate how misfolding in large RNAs can be local, and therefore also be cured locally by unfolding of small elements; this is in contrast to the function of protein chaperones, which tend to unfold the entire protein.

**Figure 1.**
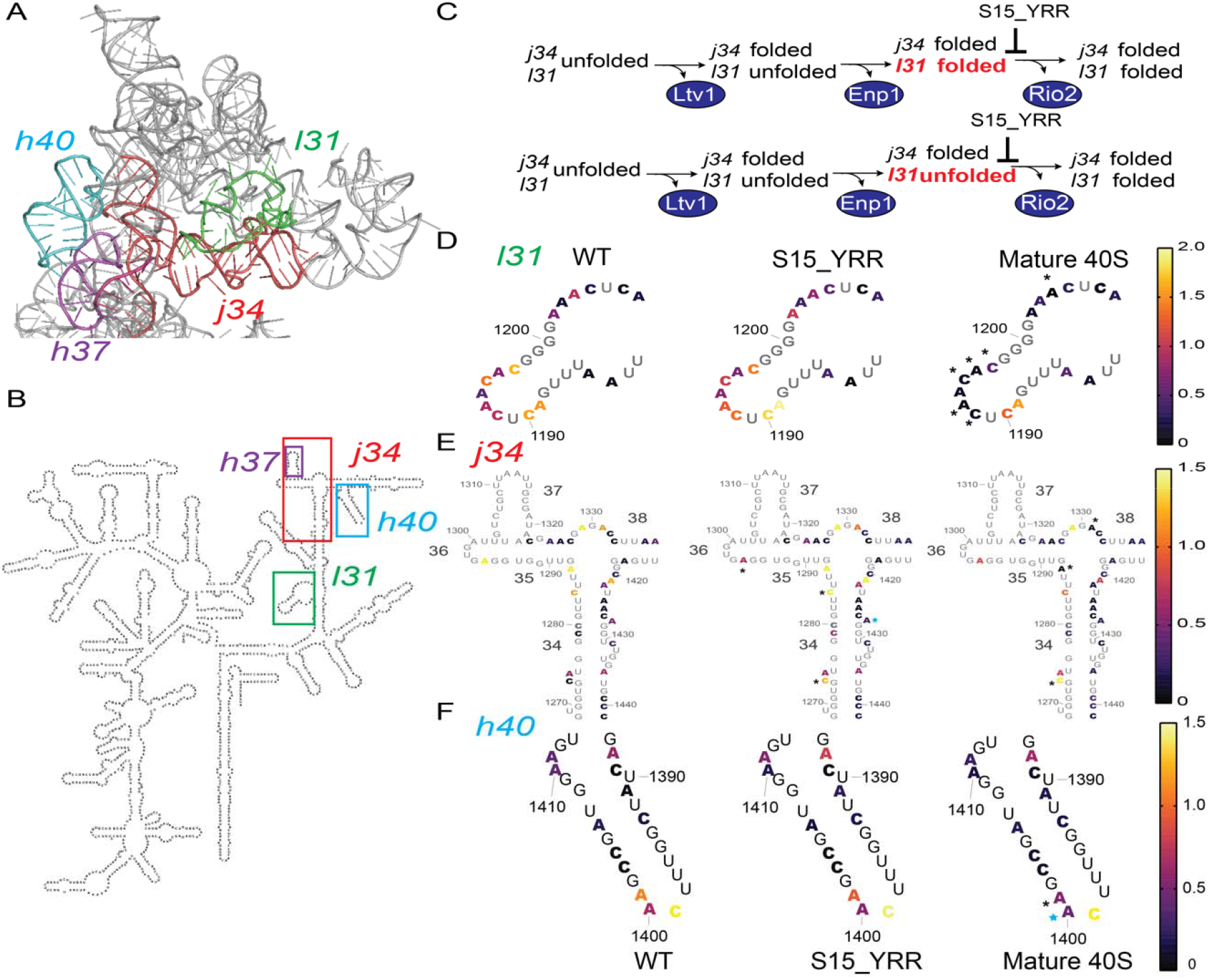
In WT yeast j34 folds before l31. A. L31 (green) and j34 (red) interact in mature 40S ribosomes (extracted from PDB ID 3J77), thereby positioning h40 (cyan) for a tertiary contact with h37 (purple). B. The locations of j34, h40 and l31 on the secondary structure of *Saccharomyces cerevisiae* 18S (from RiboVision) are highlighted with red, cyan and green boxes, respectively. C. Previous work shows the order of Ltv1, Enp1 and Rio2 release, demonstrates that j34 folds upon Ltv1 release, and that S15_YRR blocks Rio2 release (25). This mutant is used to test if l31 folds after release of Enp1 (top) or Rio2 (bottom). The two pathways make different predictions about the folding state of l31 in the S15_YRR mutant. D. DMS accessibility of nucleotides within the l31 region of 18S rRNA from WT pre-40S, S15_YRR pre-40S and mature 40S. The color of each nucleotide corresponds to its DMS accessibility as indicated in the scale bar. Changes that are statistically significant (p_adj_<0.05) using a two-way ANOVA analysis are marked with a black asterisk. n>3. E. DMS accessibility of nucleotides within the j34 region of 18S rRNA from WT pre-40S, S15_YRR pre-40S and mature 40S. Changes that are statistically significant (p_adj_<0.05) using a two-way ANOVA analysis are marked with a black asterisk. Changes that that do not rise to this statistical significance are marked with a cyan asterisk. n>3. F. DMS accessibility of nucleotides within the h40 region of 18S rRNA from WT pre-40S, S15_YRR pre-40S and mature 40S. Changes that are statistically significant (p_adj_<0.05) using a two-way ANOVA analysis are marked with a black asterisk. Changes that that do not rise to this statistical significance are marked with a cyan asterisk. n>3.

## Results

### In wild type yeast j34 folds before l31

Structural analyses have identified two late-folding RNA structural elements in the small subunit head (**Figure 1A**): the three-way junction around helices 34/35/38 (j34) and the loop at the tip of helix 31 (l31)(23, 24). The correct folding of these RNA elements (as well as the nearby binding of universally conserved ribosomal proteins) is interrogated via the ordered dissociation of the 40S assembly factors (AFs) Ltv1, Enp1 and Rio2 (**Figure 1C**), in a quality control mechanism that uses the formation of 80S-like complexes to test the ability of the nascent subunit to properly identify the start codon(25). Even though these two structural elements interact with each other in mature 40S **(Figure 1A**), structural evidence(23, 24), and our biochemical and genetic data(25) indicate that these elements form successively (**Figure S1A**). Thus, we reasoned that the successive folding of these elements, which are bound by AFs would allow us to probe how global tertiary structure is adopted by stepwise formation of individual structural elements, and what role AFs play in that process.

Structural data indicate that j34 folds before l31 (shaded path in **Figure S1A**): both j34 and l31 are unfolded in pre-40S purified *via* tagged Ltv1(23), which ensures the occupancy of Ltv1. In contrast, in a later pre-40S intermediate purified *via* tagged Nob1 (and essentially lacking Ltv1), j34 but not l31 is fully resolved(24), showing j34 can be folded without l31. These data, validated in biochemical analyses(25), also suggest that Ltv1 release triggers j34 folding. Finally, a recent structure of 80S-like ribosomes demonstrates that these lack not just Ltv1, Enp1, and Rio2, but also contain fully folded j34 and l31(26). Together, these data suggest that j34 folds before l31, making l31 the latest folding element in the small subunit head. Furthermore, the data indicate that l31 folding occurs upon either Enp1 or Rio2 release **(Figure 1C)**.

To better understand if l31 folding was delayed by an AF, we next set out to determine if l31 folds after Enp1 or after Rio2 release (**Figure 1C**). To dissect rRNA structural differences in different pre-40S intermediates we utilized DMS-MaPseq(27) (**Figure S1B)**. In brief, pre-40S intermediates were purified via Rio2-TAP to ensure the full occupancy of Rio2, and exposed to DMS, which methylates adenosines (A) or cytidines (C) that are not protected by base pairing, tertiary structure or protein binding. Modified A and C residues are misread during reverse transcription, which is assayed by deep sequencing. Therefore, by analyzing the frequency of mutations of each A and C, we can quantitatively monitor structural changes over the entire 18S rRNA. To demonstrate the effectiveness of this method and ensure that we are probing pre-40S and not the much more abundant mature 40S, we first compared the structural changes between pre-40S and mature 40S monitored by DMS-MaPseq to those observed by cryo-EM. This analysis shows that regions in pre-40S that are more accessible to DMS are located in j34, l31, helix 40 (h40), h28 and the top of h44, as expected from cryo-EM structures(23, 24, 28) **(Figure S1C)**. Those that are less accessible to DMS in pre-40S overlap the binding sites of cytoplasmic AFs **(Figure S1D)**. Moreover, analysis of the read-depth of the sequencing data demonstrates that only pre-40S samples have reads in ITS1 **(Figure S1E)**. Together, these data demonstrate that we can isolate and independently probe pre-40S subunits, and that the DMS data from these samples is fully consistent with cryo-EM structures from these intermediates.

### In wild type yeast l31 folds after Rio2 release

To dissect if l31 folds upon release of Enp1 or Rio2, we took advantage of a previously identified mutation, S15_YRR, which blocks Rio2 release after Enp1 has already dissociated (25). Thus, if l31 folds after Enp1 release (top pathway in **Figure 1C**), we would expect it to be mostly folded in the S15_YRR mutant (as in mature 40S), while l31 should be largely unfolded in the S15_YRR mutant, if l31 folds upon Rio2 release (as in WT pre-40S, bottom pathway in **Figure 1C**). Indeed, the DMS-MaPseq results show that the DMS accessibility of l31 in S15_YRR pre-40S subunits is similar to that in WT pre-40S intermediates **(Figure 1D, S1F)** strongly suggesting that l31 remains unfolded after Enp1 release, and thus folds upon Rio2 release. This finding is consistent with the location of Rio2, which is positioned to directly interact with l31(23, 24, 29) **(Figure 2A)**. In contrast to l31, which remains unfolded in S15_YRR pre-40S, two out of three significant changes in the j34 region of S15_YRR pre-40S (C1274, A1296) resemble the changes in mature 40S (**Figure 1E, S1G)**, indicating the folding of j34 in S15_YRR pre-40S, as expected based on our previous data(25). A1427 also trends towards maturation, although the change is only statistically significant in S15_YRR and not mature 40S. Release of Ltv1 and folding of j34 also leads to folding of the tip of h40, which becomes ordered in cryo-EM structures and interacts with h37, one of the helices emanating from j34(23, 24, 28) **(Figure 1A&B**). Ordering of h40 is reflected in the increased DMS protection of A1400 and A1401 in mature 40S relative to pre-40S (**Figure 1F, S1H)**. In the S15_YRR mutant this protection has been partially attained (**Figure 1F, S1H**), further supporting the interpretation that j34 is folded in the S15_YRR variant, as expected(25). The lack of full protection of j34 and h40 in the Rps15_YRR mutant is likely due to the lack of stabilization of j34 by l31.

**Figure 2.**
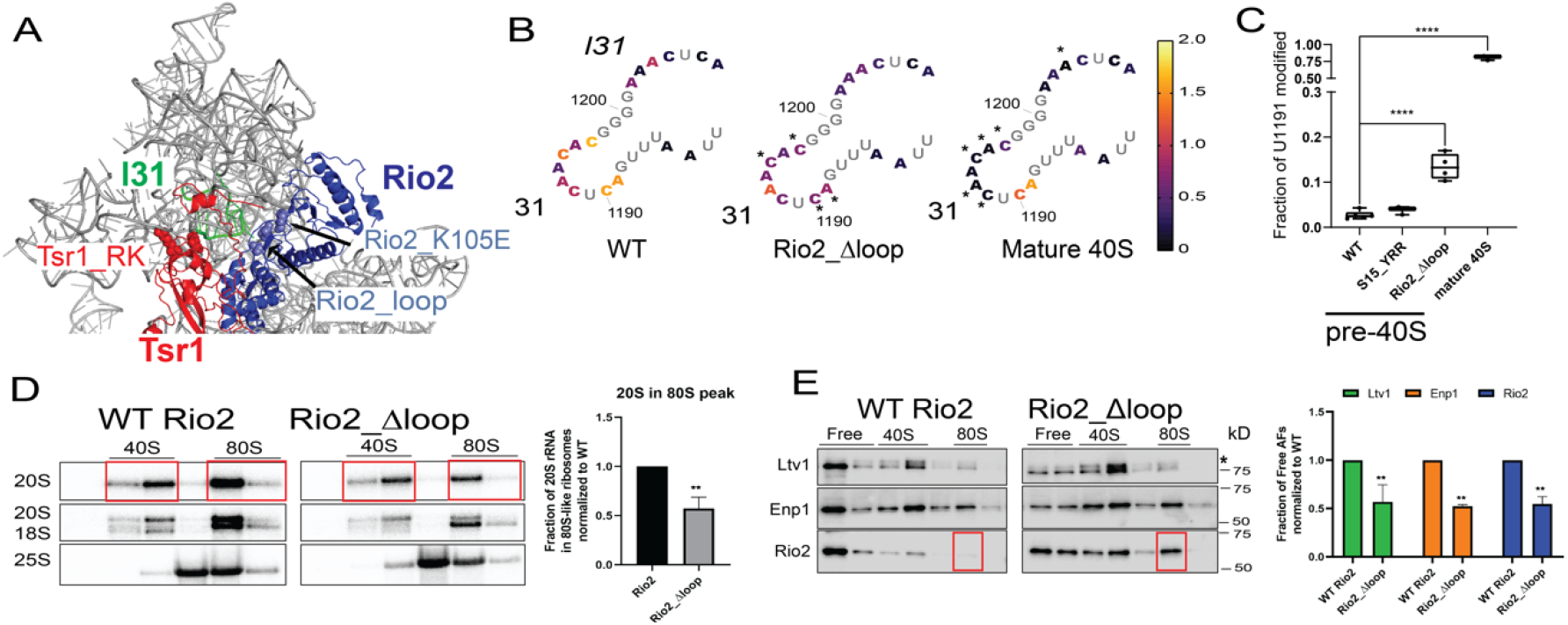
Rio2’s disordered loop keeps l31 unfolded. A. A composite structure of yeast pre-40S (PDB ID 6FAI) and mature 40S (PDB ID 3JAQ) overlaid on Rps18. Mutations in Rio2 and Tsr1 are highlighted in sphere. The loop containing the residues mutated in Rio2_loop is unresolved in all structures and the flanking residues, R129 and S145, are highlighted in sphere. Helix31 (l31) from mature 40S, which is not resolved in pre-40S, is highlighted in green. B. DMS accessibility of nucleotides within l31 of 18S rRNA from WT pre40S, Rio2_Δloop pre40S and mature 40S. The color of each nucleotide corresponds to its DMS accessibility as indicated in the scale bar. Note that the first (WT pre-40S) and third panel (mature 40S) are identical to those in Figure 1B, and shown here for direct comparison. Changes that are statistically significant (p_adj_<0.05) using a two-way ANOVA analysis are marked with a black asterisk. n>3. C. Mutational rate at U1191 from untreated WT pre40S, S15_YRR pre40S, Rio2_Δloop pre40S and mature 40S ribosomes. Significance was tested using a one-way ANOVA test. **\*\*\*\***, p<0.0001. n>3. D. Northern blots of 10-50% sucrose gradients from Fap7-depleted cells expressing plasmid-encoded wild type (WT) Rio2 or Rio2_Δloop proteins. Fap7 was depleted in glucose for over 16h to accumulate 80S-like ribosomes. The positions where 40S and 80S ribosomes sediment are indicated. (Right) Quantifications of the data on the left indicate the efficiency of formation of 80S-like ribosomes. Specifically, 20S rRNA in 80S fractions are first normalized to total 20S. This fraction of 20S in 80S-like ribosomes in Rio2_Δloop cells is further normalized to that in WT Rio2. Data are shown as mean with standard deviation. Significance was tested using an unpaired test. **\*\***, p<0.01. n=3. E. Western blots for Ltv1, Enp1 and Rio2 from the gradients in panel C. Quantifications are shown on the right. AFs in free fractions are first normalized to their total signal before normalizing the Rio2_Δloop sample to WT Rio2. Data are shown as mean with standard deviation. Significance was tested using a two-way ANOVA test. **\*\***, p<0.01. n=2.

We note that C1284 in j34 is significantly more DMS-accessible in S15_YRR pre-40S than in either WT pre-40S or mature 40S. This increased accessibility could be a reflection of the prepositioning of the j34 region for the interaction with h31. Consistent with that proposal, nearby C1279, which is adjacent to the tertiary interaction between l31 and j34, also opens up in going from WT to S15_YRR pre-40S. In summary, the DMS-MaPseq data demonstrate that j34 folds after Ltv1 release and prior to l31, consistent with previous structural(23, 24) and genetic(25) data. More importantly, the data also strongly suggest that l31 folds upon Rio2 release, immediately prior to or with the formation of 80S-like ribosomes (bottom pathway in **Figure 1C**).

### Rio2’s disordered loop keeps l31 unfolded

Above, we have shown that l31 folding follows Rio2 release, consistent with its location adjacent to l31 (**Figure 2A**). Given that helices and their loops are expected to fold rapidly during or after transcription, and not as the last element in the folding cascade, we wondered if Rio2 played a role in delaying l31 folding, until its dissociation. Intriguingly, an 18 amino acid disordered loop in Rio2 (R129-N146) is positioned to interact with l31 **(Figure 2A)**. We thus wondered if this protein loop inserts itself into l31 to prevent it from adopting its mature tertiary structure. To test this hypothesis, we deleted this loop (Rio2_Δloop(30)) and used DMS-MaPseq to probe its effects on folding of l31.

If the Rio2 loop disrupts the l31 structure, then one would expect it to adopt the folded structure - and become protected from DMS - upon deletion of the Rio2_loop. Indeed, l31 is overall less accessible to DMS in pre-40S subunits from Rio2_Δloop cells relative to WT pre-40S, and more similar to l31 from mature 40S **(Figure 2B, S1F)**. Of the five residues significantly protected in mature 40S, C1195 and C1197 are already protected in Rio2_Δloop, while C1192 is trending that way, supporting the notion that j31 folds prematurely in Rio2_Δloop cells. Notably, A1193 is more exposed in the Rio2_Δloop cells relative to mature 40S, suggesting its interaction with j34 is not yet established (see *Premature folding of l31 leads to misfolding of j34* below).

U1191 within l31 is the target of a cancer-associated amino-carboxy-propyl (acp) modification(31, 32). This modification is installed late in 40S maturation, around the time when 80S-like ribosomes form(33). l31 is folded in 80S-like ribosomes(26), suggesting that acp-modification occurs after l31 folding, and therefore can be used as an indirect read-out of l31 structure. Because the acp-modification leads to misreading of U1191 during reverse transcription(31-33), the frequency of mutations at U1191 in the non-DMS-treated sequencing libraries are a reflection of the extent of modification of that residue. These data show increased acp-modification of U1191 in Rio2_Δloop pre-40S **(Figure 2C)**, further supporting premature folding of l31 in Rio2_Δloop cells.

All studied mutants that affect l31 and j34 structure impair the formation of 80S-like ribosomes(25). Thus, we analyzed the effects from Rio2 loop deletion on the formation of 80S-like ribosomes using a previously described *in vivo* assay(25, 34-36). This assay relies on the accumulation of 80S-like ribosomes containing pre-18S rRNA (20S rRNA) and 25S rRNA in cells depleted of the ATPase Fap7, as long as 80S-like ribosomes can form (25, 34-36). Indeed, loop deletion shifts the pre-18S rRNA (20S rRNA) from 80S-like ribosomes to 40S ribosomes, under conditions that normally accumulate 80S-like ribosomes (depletion of the ATPase Fap7), indicating a deficiency in 80S-like ribosome formation **(Figure 2D)**. Next, we used Western analysis to test whether the accumulating pre-40S ribosomes retain Ltv1, Enp1 and Rio2. This analysis indicates that release of Ltv1, Enp1 and Rio2 is impaired **(Figure 2E)**, resembling the effects observed from mutation of this loop (Rio2_loop), mutation of Rio2_K105, or the nearby Tsr1 (Tsr1_RK), which also contacts h31(25) **(Figure 2A)**. Our previous analyses had demonstrated that Ltv1 release at this step requires rearrangements of h31(25). It appears that premature folding of l31 restricts these rearrangements, impairing Ltv1 release.

Most strikingly, we also observe that the nascent subunits that escape the block to form 80S-like ribosomes retain Rio2, as indicated by the accumulation of Rio2 in the 80S-like fraction **(Figure 2E**, red box). Rio2 is not normally a part of 80S-like ribosomes(26, 34). In wild type cells, l31 is only folded once 80S-like ribosomes are formed, i.e. upon Rio2 dissociation ((26) and above). The finding that Rio2 can be retained in 80S-like ribosomes when its loop is deleted, supports the model that the Rio2 loop blocks premature folding of l31. Deletion of the Rio2_Δloop allows the folding of l31 in the presence of Rio2, such that 80S-like ribosomes retain Rio2. Thus, the DMS probing and biochemical analyses provide strong evidence that Rio2’s disordered loop keeps l31 unfolded.

### Premature folding of l31 leads to misfolding of j34

We next asked why it is important for Rio2 to keep l31 unfolded until j34 folding is completed. Given that j34 forms a tertiary interaction with l31 in mature 40S **(Figure 1A)**, we hypothesized that premature formation of l31 affects folding of j34, a complex three-way junction often misfolded *in vitro(2, 3)* (**Figure 3A**). To test this hypothesis, we first scrutinized the DMS-MaPseq data of the j34 region from Rio2_Δloop pre-40S **(Figure 3B, S1G)**. C1274, A1287 and A1331 (marked with a black asterisk) change significantly towards mature 40S in the Rio2_Δloop pre-40S, although neither reaches the level of protection observed in mature 40S. In contrast, A1275, C1284, A1325 (marked with a cyan asterisk) show changes in Rio2_Δloop pre-40S that are opposite to the changes during maturation, indicating that the changes in j34 are due to misfolding instead of native folding of j34. Furthermore, A1400 and A1401 in h40 do not become protected in Rio2_Δloop intermediates **(Figure S1H)**, indicating that the interaction between helices 37 and 40 is not formed, as expected if j34 was misfolded instead of in its native structure. Lastly, misfolding (instead of natively folding) of j34 in the Rio2_Δloop intermediates is also consistent with impaired Ltv1 release in those cells **(Figure 2E**). Release of Ltv1 leads to folding of j34 (25). Thus, premature but not native folding of j34 would be expected to promote - not block - Ltv1 release.

**Figure 3.**
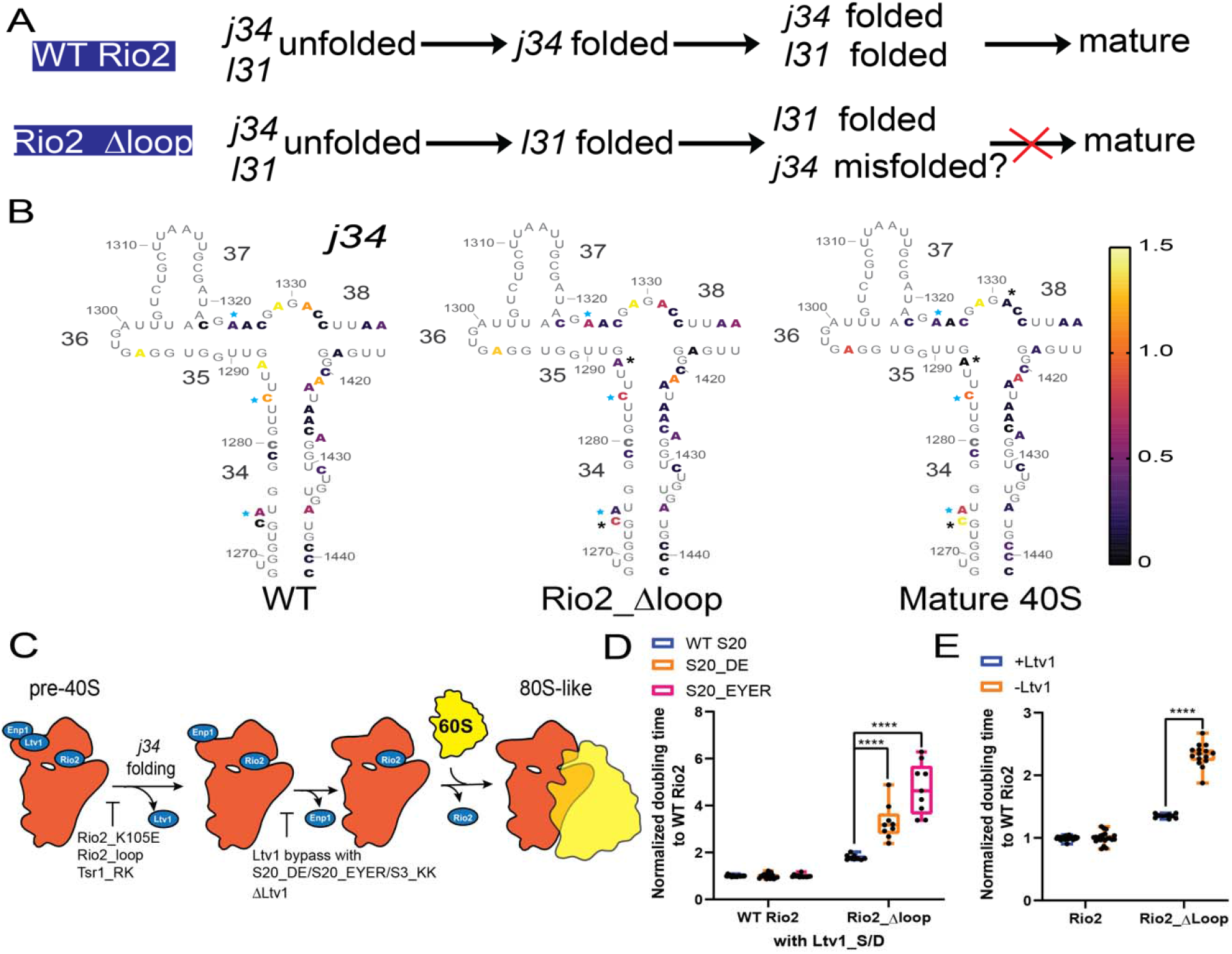
Premature folding of l31 leads to misfolding of j34. A. Model that l31 pre-mature folding traps misfolded j34 and blocks 40S maturation. B. The color of each nucleotide corresponds to its DMS accessibility as indicated in the scale bar. Note that the first (for WT pre-40S) and third panel (for mature 40S) are identical to those in Figure 1B. Changes that are statistically significant using a two-way ANOVA analysis comparing nucleotides of the entire 18S region to WT pre-40S are marked with black asterisk. *, P<0.05. Changes that are opposite in Rio2_Δloop and mature 40S are marked with a cyan asterisk. n≥3. C. Schematic release order of Ltv1, Enp1 and Rio2 prior to formation of 80S-like ribosomes. The step impaired by each mutation is indicated. D. Normalized doubling times of ΔLtv1,Gal:Rio2,Gal:S20 cells supplied with plasmids encoding Ltv1_S/D and WT Rio2 or Rio2_Δloop and WT S20 or S20_DE or S20_EYER. Significance was tested using a two-way ANOVA test. ****, p<0.0001. n≥9. E. Normalized doubling times of Gal:Rio2 or ΔLtv1,Gal:Rio2 cells supplied with plasmids encoding WT Rio2 or Rio2_Δloop. Significance was tested using a two-way ANOVA test. ****, p<0.0001. n≥10.

To further confirm that j34 is prone to misfolding in Rio2_Δloop cells, we took advantage of previously-described mutations, which destabilize j34 (S20_DE/S20_EYER with Ltv1 bypass(25)) and lead to misfolding of j34 (ΔLtv1(37)). Because these mutations destabilize the native fold of j34, they can be used as genetic probes for other elements that perturb j34 folding **(Figure 3C)**: if the premature folding of l31 in Rio2_Δloop cells leads to misfolding of j34, then we expect these mutations to amplify the effect from the Rio2_Δloop mutation and render the cells synthetically sick. Indeed, quantitative growth measurements demonstrate that the S20_DE/S20_EYER mutations **(Figure 3D)** or Ltv1 deletion **(Figure 3E)** have substantially larger effects in Rio2_Δloop cells than in wild type cells. These data demonstrate the importance of keeping l31 unfolded via the insertion of the Rio2 disordered loop to ensure proper folding of the adjacent j34.

To further test the model that l31 is kept unfolded to protect j34, we used other mutants (Tsr1_RK, Rio2_K105E, Rio2_loop) that impair 40S maturation at the same step as Rio2_Δloop(25) **(Figure 3C)**, and surround l31 **(Figure 2A)**. We combined these mutations with the molecular probes of j34 misfolding (S20_DE/S20_EYER and ΔLtv1). As expected, Tsr1_RK, Rio2_K105E, Rio2_loop also exacerbate mutations that destabilize j34 (S20_DE/S20_EYER, **Figure S3A**) or promote j34 misfolding (ΔLtv1, **Figure S3B**), producing synthetic growth defects. Moreover, these mutants are rescued by high temperature, which generally alleviates defects from misfolding **(Figure S3C)**. Finally, the growth defects of Rio2_K105E and Rio2_loop are exacerbated by low temperature **(Figure S3D)**. Together, our data indicate that Rio2 and Tsr1 cooperate to keep l31 unfolded and that pre-mature folding of l31 leads to misfolding of j34. Notably, the DMS-MaPseq data also indicate that misfolding of j34 does not lead to global changes in the small subunit structure (**Figure S2A**), although it seems that h28 starts to fold, and there may be small perturbations in the structure of h44 **(Figure S2B&C)**.

### l31 is kept unfolded by multiple factors during 40S assembly

Above we have demonstrated a role for the cytoplasmic assembly factor Rio2 in preventing the pre-mature folding of l31. However, as described above, helices and their loops are expected to fold rapidly during transcription. Thus, we wondered if additional nucleolar factors could play roles in keeping l31 unfolded early in 40S maturation. Indeed, during transcription the snoRNA snR35 binds l31 and specifies the conversion of U1191 in the loop into pseudouridine(38) **(Figure 4A)**. In a second nucleolar step, the methyltransferase Emg1 (also known as Nep1) modifies the pseudouridine to N1-methylpseudouridine. While this modification is not necessary, Emg1/Nep1 is an essential protein(39). Moreover, deletion of the nucleolar snR35 leads to accumulation of a late cytoplasmic 18S RNA precursor, 20S rRNA(38) **(Figure S4A)**. We therefore wondered if snR35 and Emg1 had a previously unappreciated role in protecting l31 from premature folding early during ribosome assembly. If this is true, then we expect deletion of these factors to block formation of 80S-like ribosomes late in assembly, when their misfolding would be detected in the previously described quality control pathway(25). To test this prediction, we used the *in vivo* assay for formation of 80S-like ribosomes described above(25, 34-36). Indeed, ΔsnR35 shifts the 20S rRNA precursor from the 80S-like fraction towards the pre-40S fraction, demonstrating that snR35 deletion impairs the formation of 80S-like ribosomes **(Figure 4B)**. Deletion of Emg1 is not viable, but can be rescued by co-deletion of the nucleolar Nop6(40). We therefore combined Gal:Emg1 with its suppressor ΔNop6 to test if Emg1 deletion affected formation of 80S-like ribosomes. To ensure that we assay only cytoplastic intermediates and not nucleolar ones, which would be unable to form 80S-like ribosomes as they have no access to 60S subunits, we used a fractionated cytoplasmic extract **(Figure S4B)**. Relative to ΔNop6 cells, depletion of Emg1 in the ΔNop6 background blocks the formation of 80S-like ribosomes **(Figure 4C)**, as expected if Emg1 has a role in keeping l31 unfolded, thereby enabling proper folding of j34. Thus, deletion of snR35 and Emg1 block the formation of 80S-like ribosomes, as expected if j34 misfolds. To further confirm that these assembly defects are linked to l31, we asked if these mutations genetically interacted with other mutations that perturb l31. Indeed, deletion of snR35 is epistatic to other mutations that perturb l31, including Rio2_Δloop, Rio2_loop, and Tsr1_RK as expected if each of these mutations has the same effect: to lead to premature folding of l31 **(Figure S4C**). We are unable to carry out this experiment for Emg1 depletion, as the combined growth defects from Gal:Emg1(ΔNop6) and Rio2_Δloop, Rio2_loop, or Tsr1_RK are too large to measure their doubling times reliably.

**Figure 4.**
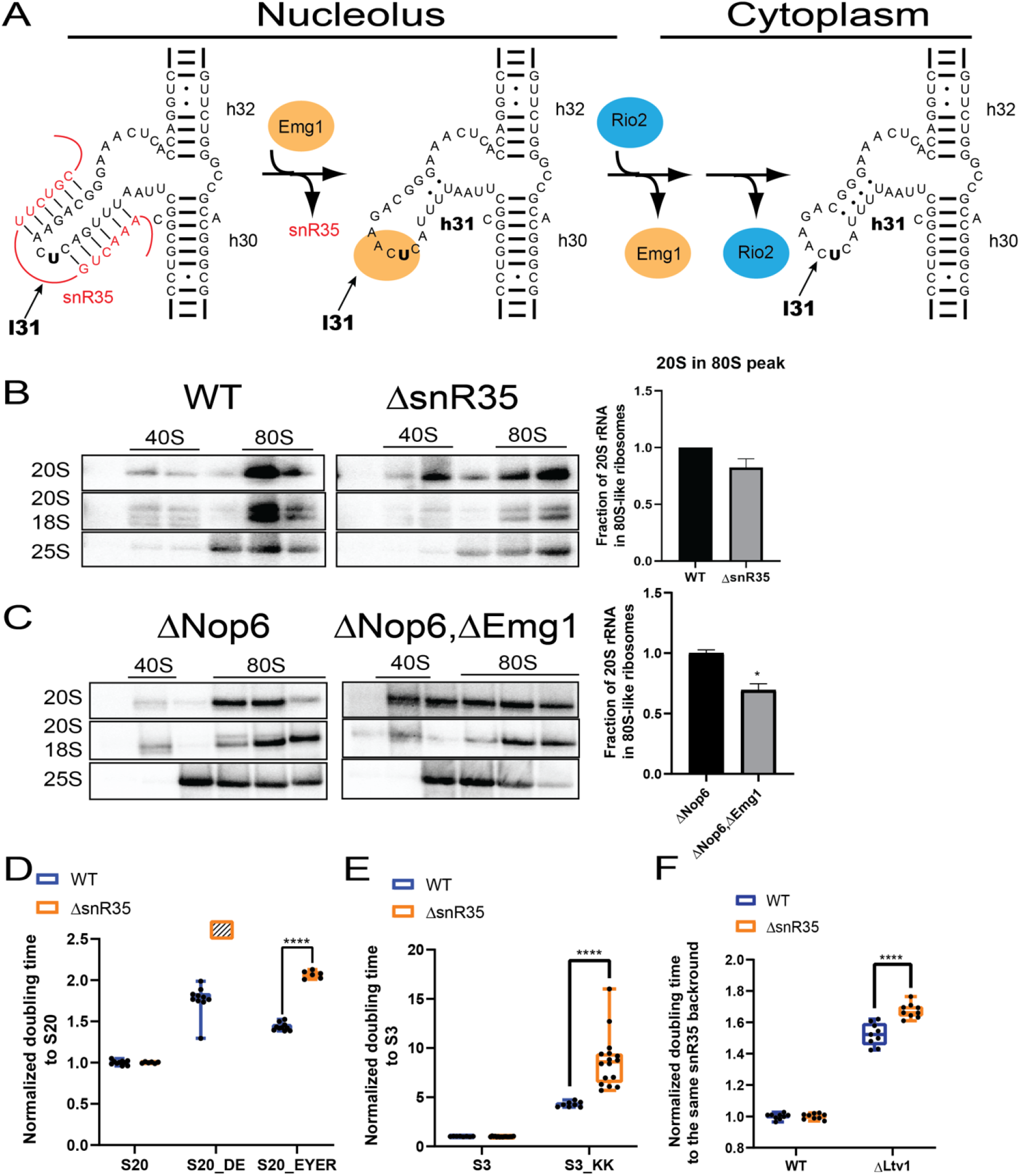
Multiple factors keep l31 unfolded during 40S assembly. A.During transcription in the nucleolus, the snoRNA snR35 binds the rRNA sequence that will become helix 31 and loop 31 (h31, l31), preventing its formation. After pseudouridylation of U1191 (in bold), snR35 dissociates, and Emg1/Nep1 binds to direct methylation of pseudo-U1191, also in the nucleolus. Emg1 is eventually replaced by Rio2, which functions in the cytoplasm. By binding the loop31 nucleotides these factors prevent the formation of its native structure. B. Northern blots of 10-50% sucrose gradients from Fap7 depleted cells with and without snR35. Fap7 was depleted in glucose for over 16h to accumulate 80S-like ribosomes. Quantifications are shown on the right. Data are shown as mean with standard deviation. n=2. C. Northern blots of 10-50% sucrose gradients from the cytoplasmic fraction of ΔNop6,Gal:Fap7 and ΔNop6,Gal:Emg1,Gal:Fap7 cells. Fap7 was depleted in glucose for over 16hrs. Quantifications are shown on the right. Data are shown as mean with standard deviation. Significance was tested using an unpaired t-test. **\***, p<0.05. n=2. D. Normalized doubling times of ΔLtv1,Gal:S20 or ΔLtv1,ΔsnR35,Gal:S20 cells supplied with plasmids encoding Ltv1_S/D and WT S20 or S20_DE or S20_EYER. Cells with Ltv1_S/D and S20_DE in ΔsnR35 do not grow, and are therefore shown as a box filled with diagonal lines. Significance was tested using a two-way ANOVA test. ****, p<0.0001. n≥6. E. Normalized doubling times of ΔLtv1,Gal:S3 or ΔLtv1,ΔsnR35,Gal:S3 cells supplied with plasmids encoding Ltv1_S/D and WT S3 or S3_KK. Significance was tested using a two-way ANOVA test. ****, p<0.0001. n≥8. F. Normalized doubling times of BY4741, ΔLtv1, ΔsnR35, ΔLtv1ΔsnR35 cells. Significance was tested using a two-way ANOVA test. ****, p<0.0001. n=9.

These data demonstrate that protection of l31 during the nucleolar stages of assembly is important for the cytoplasmic maturation of 40S ribosomes, presumably to ensure folding of j34. To test this more directly, we tested the genetic interactions between ΔsnR35 and mutants that destabilize j34 (Rps20_DE/EYER, Rps3_KK, ΔLtv1). As expected, ΔsnR35 is sensitive to the j34 destabilizing mutants **(Figure 4D-F)**, an effect that is specific to snR35 deletion and not observed upon deletion of the nearby snR57 **(Figure S4D**). Together, these data demonstrate that snR35, Emg1, Tsr1 and Rio2 function successively to prevent premature folding of loop 31 throughout assembly, thereby enabling proper folding of j34.

## Discussion

### Assembly factors act locally to isolate 3-way junctions from tertiary interactions that present kinetic traps

Like all RNAs, ribosomal RNAs are prone to misfolding *in vitro*, when assembly factors (AFs) are absent *(1-4)*. In contrast, folding *in vivo*, which is promoted by a large machinery of AFs, appears to be efficient, suggesting a role for these AFs in chaperoning RNA folding. However, the causes for RNA misfolding, and therefore the solutions that chaperones provide, have remained unclear. While the roles of alternative secondary structures in misfolding of model RNAs have been well-described(8-17), whether these play a substantial role in rRNA misfolding remains unclear. Moreover, the contributions from misfolding of helical junctions, or tertiary structures (18) remain less well-characterized, likely because it is much harder to dissect or reprogram such interactions. Finally, whether misfolding in large RNAs is local, or leads to globally altered structures also remains unclear.

Here we address these questions by probing the formation of a tertiary contact between a 3-way helical junction (j34) and the loop of a helix (l31) in the small ribosomal subunit’s P-site **(Figure 1A-B)**. Our data demonstrate that the 3-way junction is prone to misfolding, and that the misfolded intermediate becomes kinetically trapped upon formation of the tertiary interaction with the helical loop **(Figure 5)**. Moreover, the data show that a succession of AFs function throughout 40S assembly to delay the folding of the helical loop, rendering it the last element of the entire subunit to fold (**Figure 4A)**. This allows the 3-way junction to fold in isolation, and removes the kinetic trap, thereby enabling refolding of the misfolded intermediate. Thus, these chaperones act indirectly: rather than directly binding and steering the folding of the 3-way junction, they bind its interaction partners to lower the stability of the misfolded intermediate thereby allowing for its refolding. These data reveal a novel role for AFs in chaperoning rRNA folding and also demonstrate the mechanistic basis for this role.

**Figure 5.**
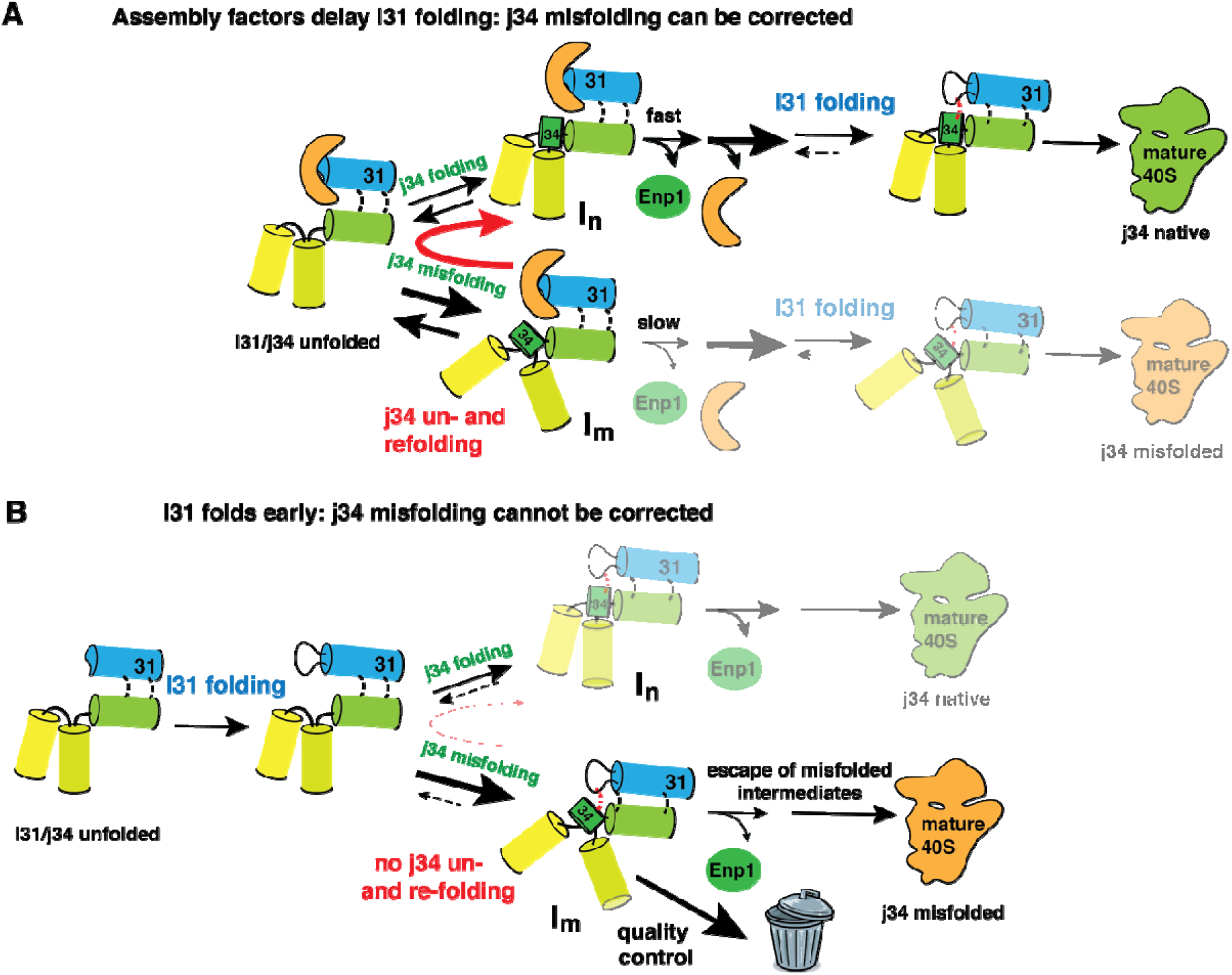
AFs choreograph rRNA folding to prevent misfolding. A. Throughout assembly, chaperones (orange) prevent the folding of l31 (shown as the loop at the end of the blue helix 31), as detailed in **Figure 4A**. In the late cytoplasmic stages shown here, Rio2 is the chaperone bound to l31, and both l31 and j34 (green disk) are initially unfolded. After dissociation of Ltv1 ((25), not shown here for clarity), j34 folds. However, during its folding, j34 partitions between folding to its native form (I_n_, top) or a misfolded intermediate (I_m_, bottom). In the presence of the chaperone, misfolding can be corrected, because the subsequent step, dissociation of Enp1, is slow when j34 is misfolded (25). This allows for unfolding of the misfolded junction, which can then re-fold (red arrow). In contrast, correct assembly (top) proceeds forward, because it is rendered irreversible via the dissociation of Enp1. Once Rio2, the chaperone (orange), also dissociates (via mechanisms detailed in (25)), l31 folds, enabling further maturation via the irreversible, eIF5B/GTP-dependent formation of 80S-like ribosomes (not shown for clarity). Thus, the chaperone assembly factors delay folding of l31, to isolate j34 and enable its proper formation by allowing refolding of its misfolded intermediate. The resulting prevalent pathway is in the foreground. B. In the absence of chaperone assembly factors, the folder order differs and l31 folds before j34 (pathway highlighted in the foreground). While the propensity of j34 to misfold is likely unchanged (as the chaperone does not bind j34, but l31), refolding of the misfolded intermediate I_m_ is no longer possible (as indicated by the dashed reverse and red arrows), as it is immediately stabilized and trapped by l31 (red interaction). Because Enp1 dissociation from this misfolded intermediate is also impaired (25), it becomes trapped and is subject to decay ((35), the quality control arrow to the trash bin). Nonetheless, Enp1 dissociates from some of the intermediates, allowing them to escape, and mature into misfolded 40S (37). Thus, the chaperone activity of Rio2 (and snR35/Enp1) is indirect: rather than directly affecting the folding of j34, they affect the stability of the intermediate to allow for its refolding.

Specifically, our data show that the snoRNA snR35 prevents premature folding of both helix 31 and its loop during transcription. After dissociation of snR35, the nucleolar Emg1 takes over in binding and chaperoning l31, followed by Rio2 (and supported by Tsr1) in the cytoplasm (**Figure 4A)**. Each of these factors binds directly to the nucleotides in l31, thereby preventing the adoption of its native structure. Thus, the l31 structure is not formed until very late in assembly, after Rio2 dissociation and just prior to formation of 80S-like assembly intermediates (26). Mechanistically, the data show that j34 (green disk in **Figure** 5) partitions between folding to a native (I_n_, top pathway in **Figure 5A**) and a misfolded form (I_m_, bottom pathway). We previously demonstrated that the step after j34 folding, dissociation of the AF Enp1, is slow when j34 is misfolded (25). This allows the misfolded I_m_ to unfold, and repartition (red arrow in **Figure 5A**). In contrast, rapid dissociation of Enp1 from natively folded I_n_ renders correct j34 folding irreversible, providing the driving force for forward assembly. Once the Rio2 chaperone (orange) dissociates from the assembling subunit, l31 (the loop at the end of the blue helix in **Figure 5)** folds, and then binds to the preformed j34. As for j34 folding, l31 formation is rendered irreversible by the eIF5B/GTP-dependent formation of 80S-like ribosomes, driving assembly forward and enabling maturation.

In the absence of the chaperone (**Figure 5B**), l31 folds prior to correct j34 folding, reflecting the rapid formation of RNA hairpin loops(41). The preformed l31 allows for the formation of the tertiary interaction between l31 and both natively and misfolded j34 (red “hooks” in **Figure 5B**), thereby “capturing” not just the correctly folded (I_n_, top pathway), but also the misfolded structure (I_m_, bottom pathway), and stabilizing it in a kinetic trap. Because misfolded *mature* ribosomes accumulate in the absence of Ltv1 (37), unfolding of j34 (dashed red line in **Figure 5B**) must now be slower than the dissociation of Enp1, thereby allowing misfolded intermediates to escape down the assembly line. Notably, the lack of substantial accumulation of pre-18S rRNA, while mature 18S rRNA is significantly reduced in the absence of Ltv1 (35), also indicates that the I_m_ intermediate is subject to degradation.

Overall, Rio2, Emg1 and snR35 function as chaperones by binding to and preventing the formation of l31, and thereby the premature formation of tertiary interactions between l31 and j34, which stabilize both natively folded and misfolded intermediates. Thus, they act by isolating the difficult-to-fold j34. However, Rip2 and Emg1 achieve this isolation not by binding the RNA element prone to misfolding, but instead by binding and blocking its interaction partner.

Notably, the ability of native tertiary interactions to stabilize not just the native structure but also misfolded intermediates has been previously only demonstrated for a small model RNA, and not in cells (18).

### Dissociation of assembly factors serves as a timer that prevents RNA misfolding

The scheme in **Figure 5B** demonstrates that escape of the misfolded intermediate is dictated by the relative speeds of RNA refolding, dissociation of Enp1, and degradation of the intermediate. The observation that misfolded mature 40S subunits accumulate in the absence of Ltv1 (37), while degradation of the I_m_ intermediate also seems to occur (35), indicates that decay of the I_m_ intermediate must occur with similar speed as Enp1 dissociation. Moreover, as described above, these findings also suggest that refolding is slow, as otherwise the misfolded mature RNA would not accumulate. Finally, this scheme makes the prediction that speeding Enp1 dissociation should lead to the accumulation of defective ribosomes, as it would favor the escape of misassembled ribosomes at the expense of quality control. Indeed, defective ribosomes accumulate in a yeast strain, where Enp1 is bound more weakly (25), thereby providing independent support for this model.

Finally, this scheme also illustrates the competing demands on ribosome assembly: while stronger binding and thereby slower dissociation of assembly factors would enhance quality control, it would also slow down assembly and promote decay of even correctly folded intermediates. Thus, AF affinities must be carefully balanced to achieve both rapid and correct ribosome assembly.

This function of dissociation of AFs in setting up a timer that regulates whether misassembled ribosomes can mature, or degraded or re-assembled, is reminiscent of the function of Prp16 and Prp22 during pre-mRNA splicing(42, 43). These splicing factors promote rearrangements of the spliceosome that lead to their inactivation if the chemical reactions do not occur rapidly enough. Conceptually analogous to what we observe here, slowing down this rejection enables mis-splicing(42, 43).

### RNA misfolding is local and corrected locally

The data also show that misfolding of j34 does not globally perturb the structure of the small subunit, but instead has local effects. This is consistent with more limited structure probing data of mature 40S from Ltv1-deficient yeast (37), as well as the structure of inactive bacterial small ribosomal subunits(44). These data support the importance of local structure in building complex RNAs through a combination of individual modules (helices) via junctions and tertiary interactions. Because complex structures are built in this combinatorial manner, RNA chaperones can work by acting locally on individual elements, instead of requiring the global unfolding of structure as is the case for protein chaperones.

### RimM chaperones j34 by binding l31 during bacterial assembly

Analogous to the roles of Rio2, Emg1 and snR35 during eukaryotic assembly, there is evidence for a role of RimM in chaperoning the correct folding of j34 in bacteria by preventing the premature formation l31 and thus its tertiary interaction with j34. When grown in the cold, bacteria lacking RimM specifically accumulate assembly intermediates with misfolded j34(45). Furthermore, these assembly intermediates are depleted of the j34-interacting uS10/uS14 (the homologs of Rps20 and Rps29)(45),(46). Additional genetic, structural and biochemical data strongly suggest that RimM chaperones j34 in bacteria by preventing the premature folding of l31, instead of binding directly to j34. First, RimM binds directly to uS19/Rps15, and docking of RimM onto pre-30S based on this interaction places it in nearly the same position as Rio2, over l31(46, 47). This position is confirmed at very high thresholding of a low-resolution EM-map(46). Moreover, mutation or depletion of RimM can be rescued by point mutations in uS13/Rps18 and uS19/Rps15 that map adjacent to l31(48). Finally, rRNA mutations that destabilize h31, or in l31 also suppress the absence or mutation of RimM(47). Together, these data indicate strongly that RimM, like Rio2 (and snR35, and Emg1) blocks the premature folding of l31, thereby chaperoning j34 by allowing its refolding instead of trapping it in a misfolded state stabilized by tertiary interactions with l31. Thus, RimM and Rio2, which have no structural similarity, have evolved independently not just to chaperone j34, but to do so via preventing the premature folding of its interaction partner l31, thereby isolating j34, and enabling its folding by removing potential kinetic traps. This convergent evolutionary solution to chaperone j34 by preventing its premature interactions with l31, suggests that isolating difficult-to-fold elements might be a wide-spread mechanism to chaperone RNA structure formation. Below, we further address this notion by analyzing misfolding of other helical junctions, and the roles of assembly factors and snoRNAs in isolating them.

### The folding of three-way junctions is a bottleneck that is choreographed by AFs

The data herein show that the 3-way junction j34 is prone to misfolding, and can be trapped in misfolded structures, when stabilized prematurely. Given the prevalence of 3-way junctions in the small ribosomal subunit RNA, where there are 7 additional ones, (**Figure S5A**), as well as in RNAs more generally, we wondered if these present common bottlenecks to folding. If so, we wanted to determine if AFs played more general roles in chaperoning their folding.

As there is no data in the literature regarding folding (or misfolding) or 18S rRNA, we interrogated the available data for folding of 16S rRNA *(1-4). In vitro* reconstitution of bacterial small ribosomal subunits occurs in three stages(4) (**Figure S5B**). An initial assembly step (stage I), where primary and secondary proteins bind to form a misfolded intermediate, termed RI, a heat-dependent refolding step to form RI* (stage II) and a final step (stage III), in which tertiary proteins bind. Probing the differential accessibility of rRNA residues to DMS and kethoxal in each stage indicates when individual regions fold. Re-inspection of these data (2, 3) indicates that there are five structural elements that misfold initially, and then refold (red in **Figure S5C**). These include the l31/j34 structure investigated herein. While both l31 and j34 adopt structure early, l31 becomes deprotected (blue in **Figure S5C**) during the heat-dependent folding step, which then allows j34 to refold upon binding of its interacting proteins uS3, uS10 and uS14 in stage III. Thus, re-analysis of previous data confirms our finding that re-folding of j34 requires l31 to be unfolded. The data herein demonstrate how the successive activities of AFs, which delay l31 folding, accomplish this task.

Similarly, there is also evidence for misfolding, unfolding and refolding of j43 (2, 3), which in mature subunits is stabilized by j34 and j39 (**Figure S5D**), and like j34/l31 is nearly identical in 16S and 18S rRNA. While the data do not provide any information about j39, j34 (mis-)folds early, and could thereby kinetically trap j43 (with or without the help of j39).

Finally, the data also suggest that j23 and j26 initially misfold, as their final folding requires the heat-dependent step (2, 3). Notably j23 is stabilized by j26 in mature subunits (**Figure S5E)**.

Thus, these examples suggest that (i) 3-way junctions are prone to misfolding *in vitro* when folding is unguided; and that (ii) misfolded 3-way structures appear to be often stabilized by interacting elements, as demonstrated for l31 herein, and indicated by our structural analysis for junctions j43 and j23. This latter observation suggests that the mechanism uncovered here for the chaperone activity of Rio2, snR35, and Emg1 - to allow the more difficult element to fold first and independent of its binding partner(s) - could prevent misfolding of RNAs more generally. We therefore next investigated the structures of assembly intermediates to ask if (i) the random and rapid folding of these junctions observed without assembly factors **(Figure S5C)** was replaced by an ordered acquisition of structures when AFs were present **(Figure S5F)**, and (ii) whether AFs might play any role in delaying the folding of structures that would otherwise be expected to fold rapidly, akin to what we demonstrate here for Rio2, snR35, and Emg1.

Indeed, analysis of the *in vivo* assembly intermediates indicates that j43 folds before j39 and j34, which stabilize j43 in mature 40S subunits **(Figure S5D&F**). Moreover, the structures(49-54) indicate that early formation of j39 is disrupted by the AFs Imp4 and Utp1. Furthermore, the observation that folding of j34 requires dissociation of the AF Ltv1 (25), indicates a role for Ltv1 in delaying the folding of j34.

Similarly, analysis of the structures of pre-40S assembly intermediates(23, 24, 50-56) indicates that j23 is formed in the earliest 40S structures that are currently available, while j26, which stabilizes j23, is formed only in the subsequent structures **(Figure S5E&F)**. This observation again suggests that native folding is promoted by stepwise acquisition of structure, where interacting partners are initially remote. Moreover, the structures also point to a role for AFs in delaying the folding of j26, which is stabilized by the early binding protein Rps23/uS12. Notably, Rps23 does not adopt its mature position, as the AFs Bms1 and Nop14 hold it in a distinct location(50-52), thereby preventing the folding of j26.

The dissection herein of the folding pathway for j34 and its interacting partner l31, as well as the above analysis of the folding pathway of three-way junctions such as j23, j26 and j43 shows that (i) misfolding of three-way junctions is common *in vitro* and in the absence of AFs, indicating that folding of 3-way junctions represents a bottleneck in the formation of native tertiary structure in rRNA, and likely other structured RNAs as well. In contrast, our analysis of the available structures of *in vivo* assembly intermediates(23, 24, 50-56) strongly suggests that (ii) *in vivo* folding of three-way junctions occurs by separating them from their interacting partners, to prevent the stabilization of misfolded intermediates and enable the conversion to the native fold; the structures and the convergent evolution of Rio2 and RimM to chaperone j34 by binding l31 also indicate that (iii) AFs can achieve this separation by sterically blocking the formation of tertiary interactions, holding interaction partners away, or blocking their folding. Thus, the role that we have demonstrated here for Rio2, Emg1 and snR35 - preventing the premature folding of l31 and its interaction with j34 to enable native folding of j34 - reflects a general role for AFs in isolating the folding of 3-way junctions. This removes tertiary interactions that kinetically trap the misfolded intermediates that appear to be common for 3-way junctions to enable native folding.

### A role for snoRNAs in modulating RNA folding

The data herein show that one role of snR35 (in addition to the pseudouridylation of U1191) is to keep l31 unfolded. Interestingly, j4, j7, j33, j34, and j39 are all bound by snoRNAs, while j23, j26 and j43 are not(57). Thus, five out of eight 3-way junctions could be chaperoned by snoRNAs. Of the remaining three, j23 and j43 fold early. Globally, most junctions include snoRNA binding sites, and moreover, 19/33 snoRNAs bind at helical junctions, indicating that chaperoning junction folding could be a previously underappreciated general role for snoRNAs, thereby helping to overcome the hurdles to native structure that we show here these junctions pose.

## Materials and Methods

### Plasmids and yeast strains

Yeast strains **(Table S1)** were generated by PCR-mediated homologous recombination(58), and confirmed by PCR, serial dilution and western blotting if antibodies were available. snoRNA deletion strains were also confirmed via Northern blotting. Plasmids **(Table S2)** were constructed using standard cloning techniques and confirmed via sequencing.

### Growth curve measurements

Cells were grown in YPD or glucose minimal media (if the supplemented plasmid is not essential) overnight, and then diluted into fresh YPD for 3-6 hours before inoculating into 96-well plates (Thermo Scientific) at a starting OD600 between 0.04 to 0.1. A Synergy.2 plate reader (BioTek) was used to record OD600 for 48 hours, while shaking at 30 °C, unless otherwise specified (37°C or 26.5°C). Doubling times were calculated using data points within the mid-log phase. Data were averaged from at least 6 biological replicates of 3 different colonies and 2 independent measurements. Statistical analyses for each measurement are detailed in the respective figure legend.

### DMS MaPseq sample preparation

Pre-40S intermediates were purified using Rio2-TAP via IgG and calmodulin beads as previously described(59). Mature 40S were purified as described(60). The purified ribosomes were treated at 30°C for 5min with 1%DMS (Sigma-Aldrich) in the presence of 80mM HEPES pH7.4, 50mM NaCl, 5mM Mg(OAc)_2_, 0.2uM RNaseP RNA. DMS reactions were stopped by addition of 0.4-vol buffer (1M β-ME, 1.5M NaOAc, pH 5.2) and purified using phenol chloroform precipitation.

### DMS MaPseq RNA library preparation

Generation of the RNA sequencing library was adapted from a previous protocol(61). RNAs were fragmented with 20mM MgCl_2_ at 94°C for 10min, before separation on a denaturing 15% gel. RNA fragments between 50-80nt were cut out and eluted overnight in RNA gel extraction buffer (0.3M NaOAc, pH5.5, 1mM EDTA, 0.25% v/v SDS). RNA fragments were dephosphorylated using T4 PNK (NEB), ligated to the adaptor using T4 RNA ligase II truncated (NEB) and gel purified. Reverse transcription was carried out with TGIRT III (InGex) in 50mM Tris HCl, pH 8.3, 75mM KCl, 3mM MgCl2, 1mM dNTPs, 5mM DTT, 10U SUPERase·In (Invitrogen), for 1.5hrs at 60°C. RNAs were hydrolyzed by addition of 250mM NaOH and the cDNAs were circularized using CircLigase II (Lucigen). The circularized cDNAs were sent to Genomics Core at Scripps Florida for deep sequencing using either single-end or pair-end Illumina NextSeq 500 platform.

### DMS MaPseq data processing

Processing of the DMS MaPseq data followed the pipeline from GitHub code (https://github.com/borisz264/mod_seq). Briefly, raw sequencing reads were first trimmed of their adaptor sequences via Skewer(62) (Adaptor sequences: AGATCGGAAGAGCACACGTCTGAACTCCAGTCA for single-end sequencing reads and R1: AGATCGGAAGAGCACACGTCTGAACTCCAGTCA, R2: AGATCGGAAGAGCGTCGTGTAGGGAAAGAGTGT for pair-end sequencing reads.) Shapemapper 2.0(63) was used to trim the first and last 5nt of each read as well as low-quality nucleotides, align reads to *S. cerevisiae* 20S rRNA (Saccharomyces Genome Database), and count mutations and read coverage at each rRNA position. Raw sequencing data are deposited in GEO database under accession number **GSE160161**.

### *DMS MaPseq* data analysis

The first 5 and last 5nt of 20S rRNA as well as nucleotides 1766-1785 were excluded because of their low read depth. Native 18S rRNA modifications sites were analyzed separately. We first separately averaged the mutational rates of A, C, G and U in each sample (DMS-treated and untreated, and each mutant). This analysis confirmed that DMS only increases the mutational rate of A and C, as expected because only DMS-modifications at A and C lead to mutations during reverse transcription. All subsequent analyses were thus done only with A and C. To calculate the DMS accessibility for each nucleotide, we divided that nucleotide’s mutational rate by the average mutational rate of all untreated A (or C) nucleotides, before subtracting the value for the untreated sample from the DMS-treated sample.

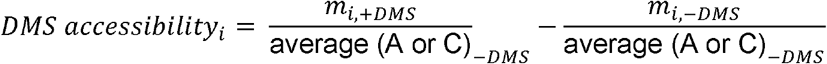

Where m_i_ is the mutational rate of nucleotide i, either with (+DMS) or without (-DMS) DMS. At least three biological replicates were collected for each sample. Raw mutational rates are summarized in file **“Summary of normalized DMS accessibility**.**txt”** under **GSE160161**.

### In vivo subunit joining assay

Cells were grown to mid-log phase in galactose before inoculating into YPD for at least 16 hours to deplete Fap7. Formation of 80S-like ribosomes was assayed by sucrose gradient fractionation and Northern blotting as previously described(25, 34-36), and the fraction of 20S rRNA in 80S-like ribosomes quantified. Dissociation of assembly factors was assayed by Western blotting, and the fraction of each AF in pre-40S ribosomes quantified. Cellular fractionation for ΔNop6,Gal:Emg1 cells was carried out exactly as previously described(35). At least two biological replicates were assayed for each mutant.

### Quantification and statistical analysis

Western blots were imaged using the ChemiDoc MP Imaging System from Biorad after applying luminescence substrates (Invitrogen). Northern blots were visualized by Typhoon™ FLA 7000(GE Healthcare). Image Lab (Biorad) and Quantity One (Biorad) were used to analyze Western blots and Northern blots, respectively.

Statistical analyses were performed using GraphPad Prism version 8.3 (GraphPad Software, San Diego, California). Statistical tests were used as indicated in the respective figure legends.

## Acknowledgments

We thank Dr. Karunadharma from the Scripps Florida Genomics Core for library preparation and sequencing, Z. Zhao for technical support with the running and debugging of Github codes and D. Herschlag and members of the Karbstein laboratory for discussion and comments on the manuscript.

This work was supported by NIH grants R01-GM117093, R01-GM086451, R35-GM136323, and HHMI Faculty Scholar grant 55108536 to K.K., and a Farris Foundation fellowship to H.H..

## Notes

**Competing Interest Statement:** No competing interests.

### Competing Interest Statement

The authors have declared no competing interest.

### Summary of Updates

The presentation of the data has been significantly changed.

