## supplemental data for "Assembly factors chaperone rRNA folding by isolating helical junctions that are prone to misfolding"

* Katrin Karbstein.

**This PDF file includes:**

Figures S1 to S5

Tables S1 to S4

SI References

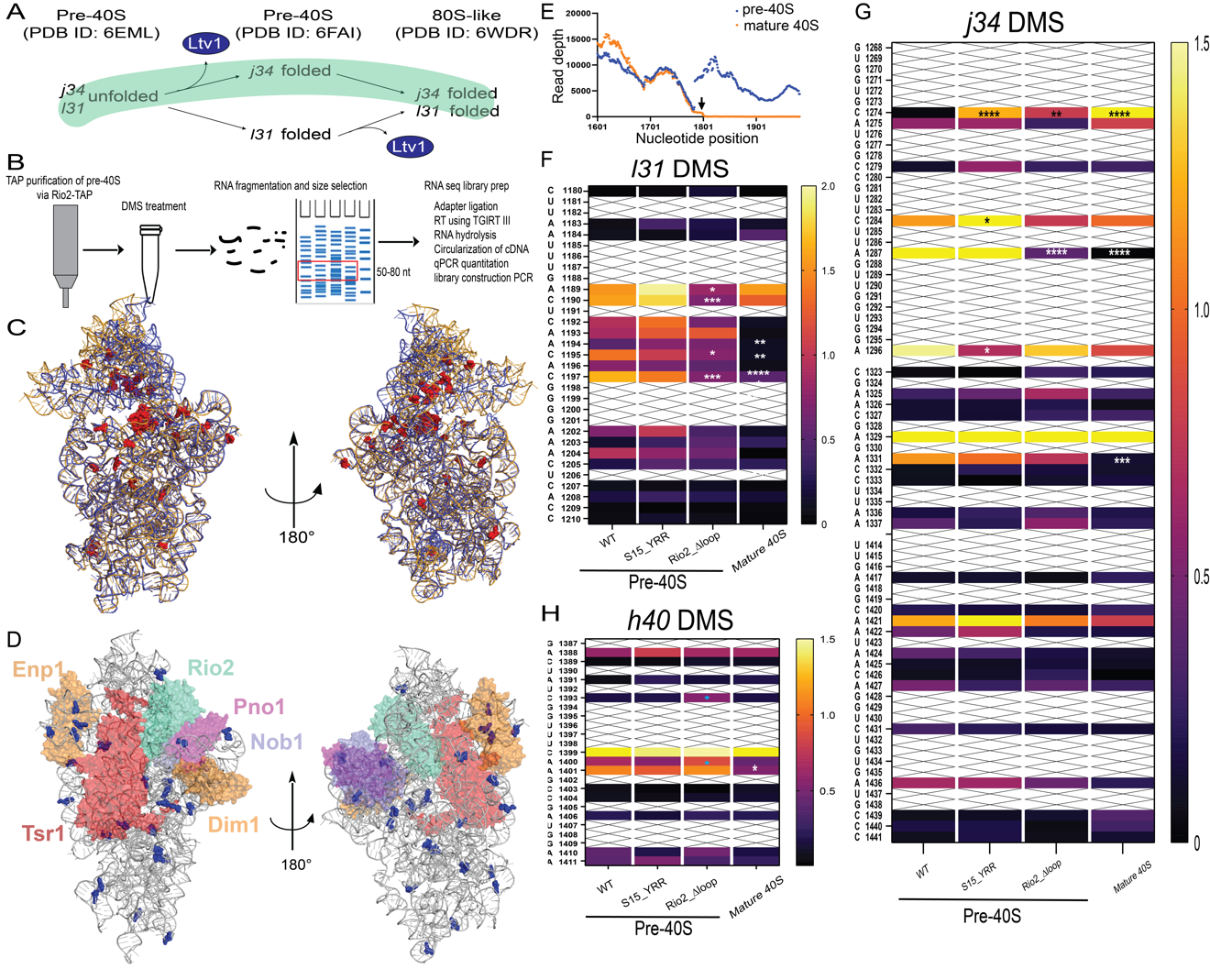

**Figure S1. DMS-MaPseq isolates pre-40S structure**

A. Two possible pathways for folding of j34 and l31 during 40S maturation. The green-shaded pathway is taken in cells.
B. Schematic outline of the DMS-MaPseq protocol to quantify structural differences of rRNA in different 40S intermediates.
C. Differential DMS accessibility between WT pre-40S and mature 40S visualized on 3D structures. 18S rRNAs from WT pre-40S(3) (in orange PDB ID 6FAI) and mature 40S(5) (in blue PDB ID 3J77) are aligned on the body. Nucleotides that are significantly more accessible in WT pre-40S are highlighted in red spheres. Significance was tested using a two-way ANOVA test. n$\geq$3.
D. A composite pre-40S structure aligning pre-40S from yeast(3, 7, 8) (PDB ID 6FAI for Tsr1, Rio2, Enp1, Pno1, and PDB ID 6RBD for Dim1) and human^3^ (PDB ID 6G18 for Nob1) based on Rps15. Nucleotides that are significantly less accessible in WT pre-40S are highlighted in blue spheres. Significance was tested using a two-way ANOVA test. n$\geq$3.
E. Read depth of typical DMS-MaPseq reads from WT pre-40S and mature 40S. The end of 18S rRNA is marked with black arrow.
F. Heat map of DMS MaPseq results of l31 of 18S rRNA from WT pre-40S, S15_YRR pre-40S, Rio2_Δloop pre-40S and mature 40S. Mean of DMS accessibility is shown for each cell type at each nucleotide. DMS accessibility larger than 2.0 is colored in yellow. G and U are not DMS-modified and thus shown as box with X. Changes that are statistically significant using a two-way ANOVA analysis are marked with an asterisk. *, P_adj_<0.05. **, P_adj_<0.01.***, P_adj_<0.001. ****, P_adj_<0.0001. n≥3.
G. Heat map of DMS MaPseq results of the j34 region of 18S rRNA from WT pre-40S, S15_YRR pre-40S, Rio2_Δloop pre-40S and mature 40S. DMS accessibility larger than 1.5 is colored in yellow. Changes that are statistically significant using a two-way ANOVA analysis are marked with an asterisk. *, P_adj_<0.05. **, P_adj_<0.01.***, P_adj_<0.001.

****, P_adj_<0.0001. n≥3.

H. Heat map of DMS MaPseq results of h40 of 18S rRNA from WT pre-40S, S15_YRR pre-40S, Rio2_Δloop pre-40S and mature 40S. DMS accessibility larger than 2.0 is colored in yellow.

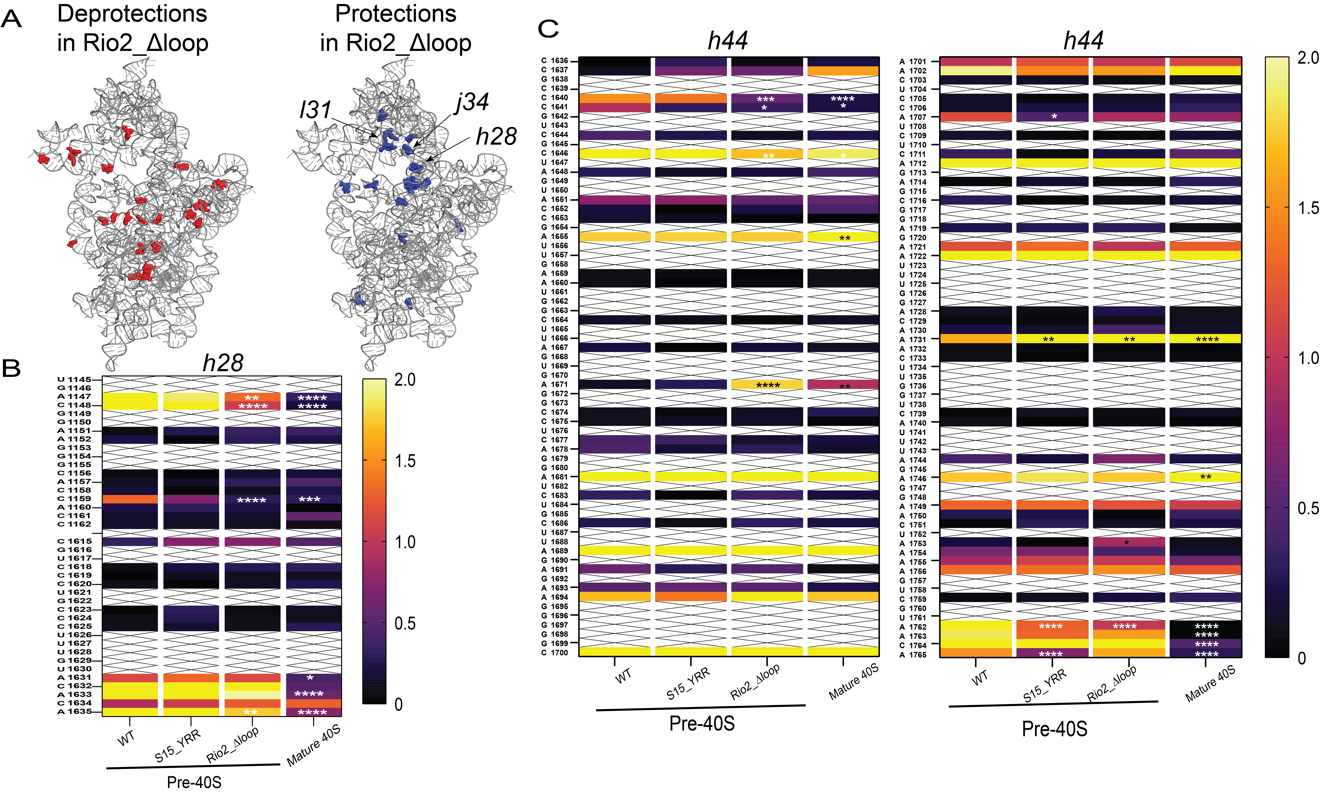
**Figure S2. The Rio2_Δloop variant induces changes in h28 and h44**

A. The Rio2_Δloop mutation does not globally perturb 40S structure. Shown here are statistically significant deprotections and protections mapped on the subunit relative to wild type pre-40S. Most novel protections arise in l31, and j34, as well as h28 and are described herein. Most deprotections are similar to those observed in mature 40S (**Figure S1C**), suggesting partial loss of assembly factors during purification with the Rio2_Δloop deletion. Significance was tested using a two-way ANOVA test. n$\geq$3.
B. Heat map of DMS MaPseq results of h28 of 18S rRNA from WT pre-40S, S15_YRR pre-40S, Rio2_Δloop pre-40S and mature 40S. DMS accessibility larger than 2.0 is colored in yellow. Changes that are statistically significant using a two-way ANOVA analysis are marked with an asterisk. P_adj_<0.05. **, P_adj_<0.01.***, P_adj_<0.001. ****, P_adj_<0.0001. n≥3.
C. Heat map of DMS MaPseq results of h44 of 18S rRNA from WT pre-40S, S15_YRR pre-40S, Rio2_Δloop pre-40S and mature 40S. DMS accessibility larger than 2.0 is colored in yellow. Changes that are statistically significant using a two-way ANOVA analysis are marked with an asterisk. P_adj_<0.05. **, P_adj_<0.01.***, P_adj_<0.001. ****, P_adj_<0.0001. n≥3.

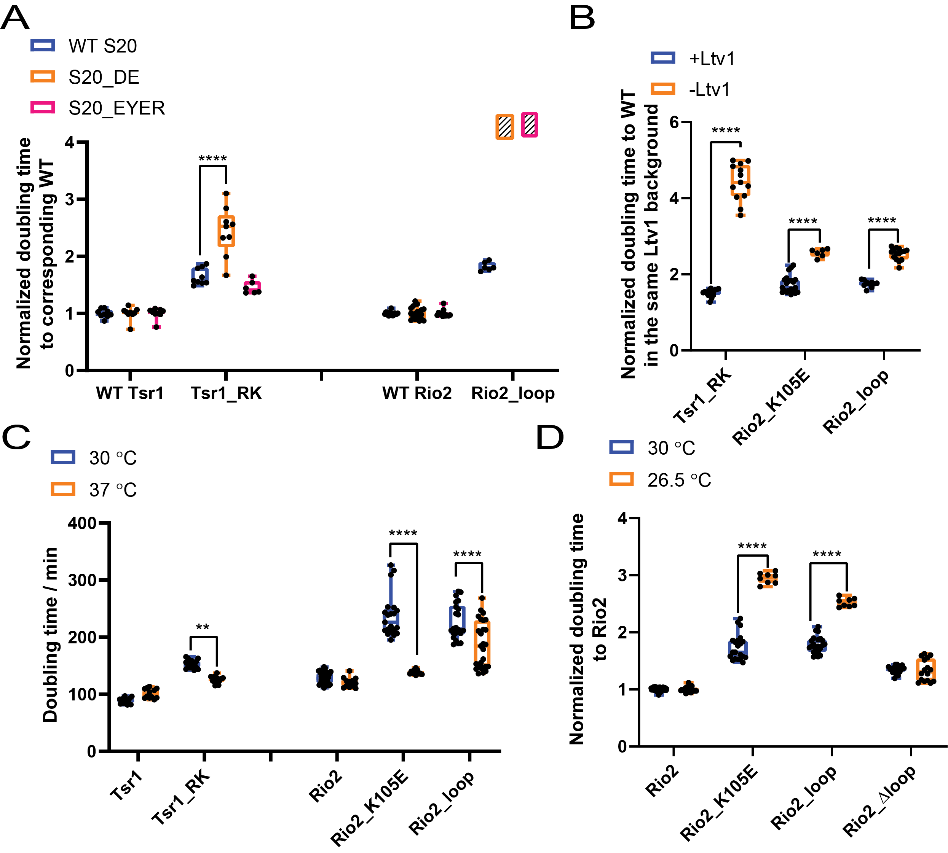

**Figure S3.**

A. Normalized doubling times of ΔLtv1,Gal:Tsr1,Gal:S20 or ΔLtv1,Gal:Rio2,Gal:S20 cells supplied with plasmids encoding Ltv1_S/D and WT Tsr1 or Tsr1_RK, or WT Rio2 or Rio2_loop and WT S20 or S20_DE or S20_EYER. Cells with Ltv1_S/D and S20_DE or S20_EYER and Rio2_loop do not grow and are therefore shown as a box filled with diagonal lines. Significance was tested using a two-way ANOVA test. ****, P<0.0001. n≥6.
B. Normalized doubling times of yeast cells encoding Tsr1_RK, Rio2_K105E and Rio2_loop in the presence and absence of Ltv1. Significance was tested using a two-way ANOVA test. ****, P<0.0001. n≥6.
C. Doubling times of Gal:Tsr1 cells supplemented with plasmids encoding WT Tsr1 or Tsr1_RK measured at 30 °C or 37 °C. Doubling times of Gal:Rio2 cells supplemented with WT Rio2 or Rio2_K105E or Rio2_loop plasmid measured at 30 °C or 37 °C. Significance was tested using a two-way ANOVA test. ****, P<0.0001. n≥12.
D. Normalized doubling times of Gal:Rio2 cells supplemented with plasmids encoding WT Rio2 or Rio2_K105E or Rio2_loop or Rio2_Δloop measured at 30 °C or 26.5 °C. Significance was tested using a two-way ANOVA test. ****, P<0.0001. n≥8.

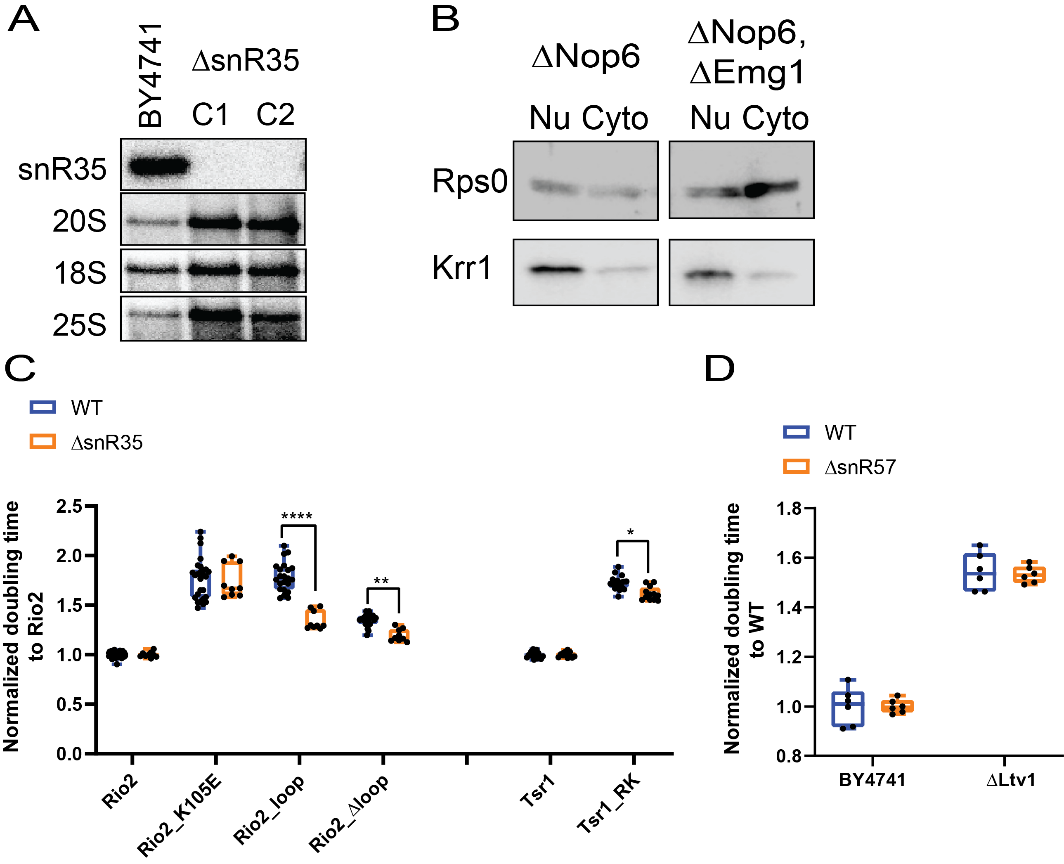

**Figure S4. snR35 deletion leads to j34 misfolding**

A. Northern blots of total RNA from BY4741 or ΔsnR35 cells. snR35, 20S, 18S and 25S are probed.
B. Western blots for Rps0(cytoplasmic) and Krr1 (nuclear) of nuclear and cytoplasmic fractions in Figure 4C.
C. Normalized doubling times of cells supplemented plasmids encoding with WT Rio2, Rio2_K105E, Rio2_loop or Rio2_Δloop in the presence (Gal:Rio2) and absence (ΔsnR35,Gal:Rio2) of snR35. Normalized doubling times of cells supplemented plasmids encoding with WT Tsr1 or Tsr1_RK in the presence (Gal:Tsr1) and absence (ΔsnR35,Gal:Tsr1) of snR35. Significance was tested using a two-way ANOVA test. **, P<0.01. ****, P<0.0001. n≥9.
D. Normalized doubling times of BY4741, ΔLtv1, ΔsnR57, ΔLtv1ΔsnR57 cells. n=6.

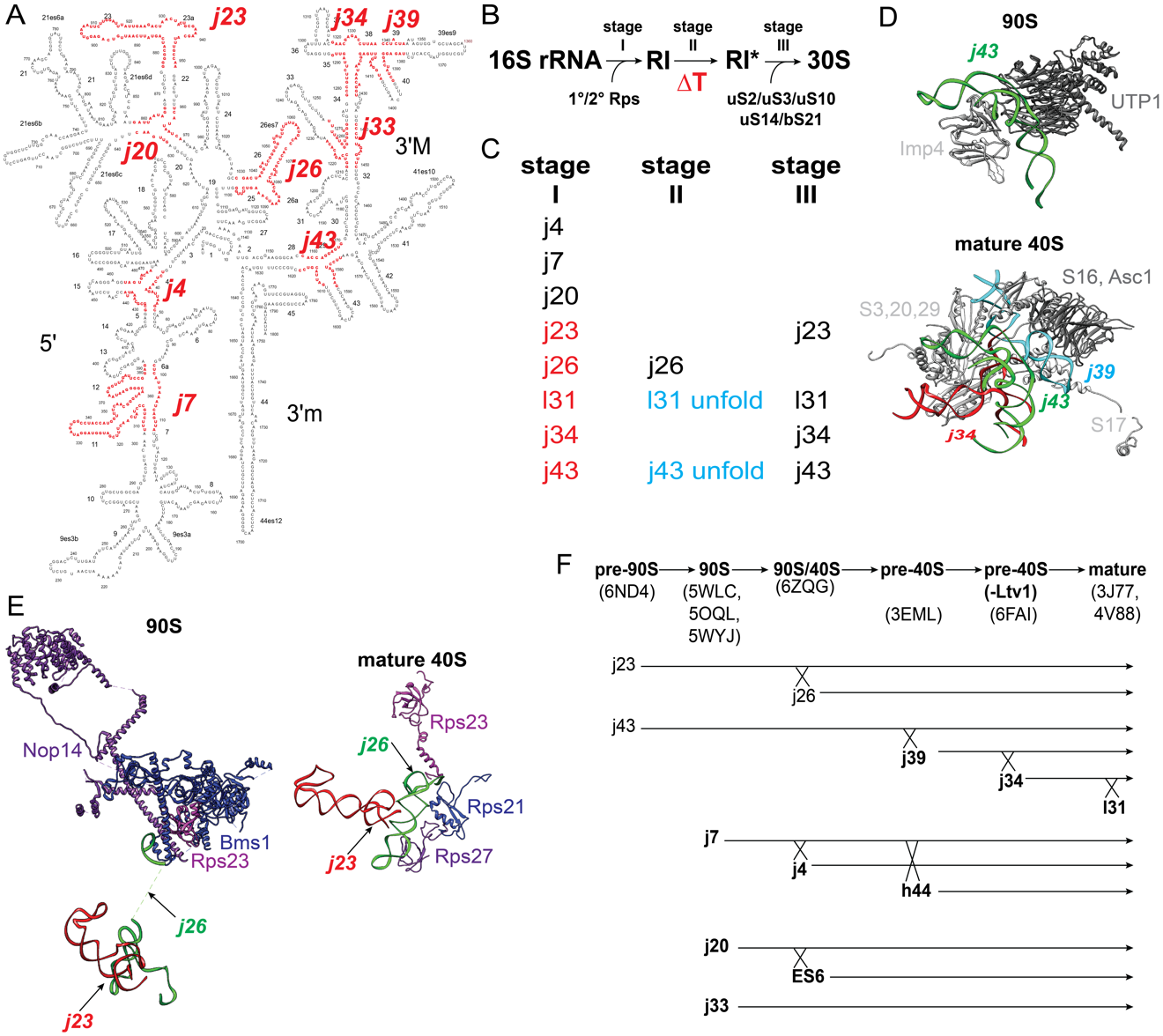

**Figure S5. 3-Way junctions in rRNA Folding.**

A. Secondary structure of 18S rRNA from Saccharomyces cerevisiae (RiboVision) with j4, j7, j20, j23, j26, j33, j34, j39, j43 highlighted in red.
B. Three stages of bacterial small ribosomal subunit assembly during *in vitro* reconstitution.
C. Folding status of indicated RNA elements with folded junctions in black, misfolded RNA in red and unfolded elements in cyan.
D. Visualizing j43 (in green) on 90S(6) (PDB ID. 5WYJ) together with Imp4 (in light gray) and UTP1 (in dark gray). Visualizing j43 (in green) on composite mature 40S(5) (PDB ID. 3J77 and 4V88) together with S3,20,29,17 (in light gray) and S16, Asc1 (in dark gray).
E. Visualizing j23 (in red) and j26 (in green) on 90S(4) (PDB ID. 5WLC) and mature 40S(5) (PDB ID 3J77). Nop14 (in purple), Bms1 (in blue) and Rps23 (in magenta) are shown in the 90S structure (left). Rps21 (in blue), Rps23 (in magenta) and Rps27 (in purple) are shown in the mature 40S structure (right).
F. Folding status of rRNA junctions in different 40S assembly intermediates. The PDB IDs for each intermediate are listed (3-6, 9-11). Only folded junctions are depicted and those captured by snoRNAs during transcription are shown in bold.

**Table S1: Yeast strains used in this study**

| **Strains** | **Description** | **Genotype** | **Reference** |
| --- | --- | --- | --- |
| YKK200 | BY4741 | MATa His3-1 Leu2-0 Met15-0 Ura3-0 |  |
| YKK87 | Rio2-TAP | BY4741, Rio2-TAP(HIS) | (1) |
| YKK1181 | Gal:Rio2 | BY4741, Gal:Rio2(KAN) | (2) |
| YKK1183 | Gal:Rio2,Gal:S15 | BY4741, Gal:Rio2(KAN), Gal:S15(NAT) | (2) |
| YKK1281 | Gal:Rio2,Gal:Fap7 | BY4741, Gal:Rio2(KAN), Gal:Fap7(HYG) | (2) |
| YKK1390 | ΔLtv1,Gal:Rio2,Gal:S20 | BY4741, Gal:S20(HYG), Gal:Rio2(NAT), Ltv1::KAN | This work |
| YKK1190 | ΔLtv1,Gal:Rio2 | BY4741, Gal:Rio2(KAN), Ltv1::HYG | (2) |
| YKK218 | Gal:Fap7 | BY4741, Gal:Fap7(KAN) | (1) |
| YKK1336 | ΔsnR35,Gal:Fap7 | BY4741, snR35::KAN,Gal:Fap7(HYG) | This work |
| YKK1435 | ΔNop6,Gal:Fap7 | BY4741, Nop6::KAN, Gal:Fap7(HYG) | This work |
| YKK1434 | ΔNop6,Gal:Emg1,Gal:Fap7 | BY4741, Gal:Emg1(NAT), Nop6::KAN, Gal:Fap7(HYG) | This work |
| YKK729 | ΔLtv1,Gal:S20 | BY4741, Ltv1::KAN, Gal:Rps20 (HYG) | (12) |
| YKK1400 | ΔLtv1,ΔsnR35,Gal:S20 | BY4741, snR35::KAN, Ltv1::NAT, Gal:S20(HYG) | This work |
| YKK73 | ΔLtv1 | BY4741, Ltv1::KAN | (1) |
| YKK1333 | ΔsnR35 | BY4741, snR35::KAN | This work |
| YKK1389 | ΔLtv1,ΔsnR35 | BY4741, snR35::KAN, Ltv1::NAT | This work |
| YKK762 | ΔLtv1,Gal:S3 | BY4741, Ltv1::KAN, Gal:Rps3 (NAT) | (12) |
| YKK1395 | ΔLtv1,ΔsnR35,Gal:S3 | BY4741, snR35::KAN, Ltv1::NAT, Gal:S3(HYG) | This work |
| YKK1436 | ΔLtv1,Gal:Tsr1,Gal:S20 | BY4741, Ltv1::KAN, Gal:Tsr1(NAT), Gal:S20(HYG) | This work |
| YKK642 | ΔLtv1,Gal:Tsr1 | BY4741, Ltv1::Kan; Gal:Tsr1 (NAT) | This work |
| YKK1149 | Gal:Tsr1 | BY4741, Gal:Tsr1(KAN) | (1) |
| YKK1423 | ΔsnR35,Gal:Rio2 | BY4741, snR35::KAN,Gal:Rio2(NAT) | This work |
| YKK1420 | ΔsnR57 | BY4741, snR57::KAN | This work |
| YKK1438 | ΔsnR57, ΔLtv1 | BY4741, snR57::KAN, Ltv1::HYG | This work |
| YKK1448 | ΔsnR35,Gal:Tsr1 | BY4741, snR35::KAN,Gal:Tsr1(NAT) | This work |

**Table S2: Plasmids used in this study**

| **Plasmid** | **Description** | **Vector** | **Reference** | **Detailed description** |
| --- | --- | --- | --- | --- |
| PKK3644 | Rio2-TAP | TEF 416 | This work | C terminal TAP |
| PKK30457 | Rio2_Δloop-TAP | TEF416 | This work | Deleting Rio2 loop(129-146) |
| PKK30216 | S15_YRR | TEF415 | (2) | Rps15_Y123I,R127R130E |
| PKK3350 | WT Rio2 | TEF413 | (13) |  |
| PKK30354 | Rio2_Δloop | TEF413 |  | Deleting Aa from 129 to 146. |
| PKK30208 | WT Rio2 | Cyc1 415 |  |  |
| PKK30369 | Rio2_Δloop | Cyc1 415 |  | See PKK30354 |
| PKK3557 | Ltv1_S/D | TEF 416 | (14) | Ltv1_S336D,S339D,S342D |
| PKK3607 | Ltv1_S/D | TEF413 |  | See PKK3557 |
| PKK3890 | WT S20 | TEF415 | (12) |  |
| PKK3934 | S20_DE | TEF415 |  | D113K,E115K |
| PKK3891 | S20_EYER | TEF415 |  | E80K,Y82A,E83K,R85E |
| PKK3295 | WT Tsr1 | TEF416 | (2) |  |
| PKK3716 | Tsr1_RK | TEF416 |  | R709E,K712E |
| PKK30183 | WT Tsr1 | TEF415 |  |  |
| PKK30184 | Tsr1_RK | TEF415 |  | See PKK3716 |
| PKK30356 | Rio2_loop | TEF413 | (2, 13) | R129A, H133A, R136A, R139A, D140A, K143A, K144A; see also (13) |
| PKK30371 | Rio2_loop | Cyc1 415 |  |  |
| PKK3795 | Rio2_K105E | TEF413 |  |  |
| PKK30211 | Rio2_K105E | Cyc1 415 |  |  |
| PKK3521 | WT S3 | TEF416 | (12) |  |
| PKK3933 | S3_KK | TEF416 |  | S3_K7D,K10E |

**Table S3: Oligos used in this study**

| **Description** | **Sequence** | **Reference** |
| --- | --- | --- |
| 18S probe | TCCCCTAACTTTCGTTCTTG | (15) |
| 20S probe | GCT CTC ATG CTC TTG CC |  |
| 25S probe | GCCCGTTCCCTTGGCTGTG |  |
| snR35 probe | TGATGATCTCTCCGATGGACTTGACGC | (16) |
| snR57 probe | GAAGAATTCCTAATTCACAATATG | This work |

**Table S4: Yeast strains and plasmids used in each figure**

| **Figure** | **Yeast strains** | **Plasmids** | **Vectors** |
| --- | --- | --- | --- |
| DMS-MaPseq results | BY4741 | / | / |
|  | Rio2-TAP | / | / |
|  | Gal:Rio2 | Rio2_Δloop-TAP | TEF416 |
|  | Gal:Rio2,Gal:S15 | Rio2-TAP+S15_YRR | TEF416 for Rio2 plasmid and TEF 415 for S15 plasmid |
| Figure 2C, 2D | Gal:Rio2,Gal:Fap7 | Rio2 | Cyc1 415 |
|  |  | Rio2_Δloop |  |
| Figure 3D | ΔLtv1,Gal:Rio2,Gal:S20 | Ltv1_S/D + WT S20+ WT Rio2 | TEF416 for Ltv1_S/D, TEF 413 for Rio2 plasmids and TEF 415 for S20 plasmids |
|  |  | Ltv1_S/D + S20_DE + WT Rio2 |  |
|  |  | Ltv1_S/D + S20_EYER + WT Rio2 |  |
|  |  | Ltv1_S/D + WT S20+ Rio2_Δloop |  |
|  |  | Ltv1_S/D + S20_DE + Rio2_Δloop |  |
|  |  | Ltv1_S/D + S20_EYER + Rio2_Δloop |  |
| Figure 3E | Gal:Rio2 | Rio2 | Cyc1 415 |
|  |  | Rio2_Δloop |  |
|  | ΔLtv1,Gal:Rio2 | Rio2 |  |
|  |  | Rio2_Δloop |  |
| Figure 4B | Gal:Fap7 | / | / |
|  | ΔsnR35,Gal:Fap7 | / | / |
| Figure 4C, S4B | ΔNop6,Gal:Fap7 | / | / |
|  | ΔNop6,Gal:Emg1,Gal:Fap7 | / | / |
| Figure 4D | ΔLtv1,Gal:S20 | Ltv1_S/D + WT S20 | TEF416 for Ltv1_S/D and TEF 415 for S20 plasmids |
|  |  | Ltv1_S/D + S20_DE |  |
|  |  | Ltv1_S/D + S20_EYER |  |
|  | ΔLtv1, ΔsnR35, Gal:S20 | Ltv1_S/D + WT S20 |  |
|  |  | Ltv1_S/D + S20_DE |  |
|  |  | Ltv1_S/D + S20_EYER |  |
| Figure 4E | BY4741 | / | / |
|  | ΔLtv1 | / | / |
|  | ΔsnR35 | / | / |
|  | ΔLtv1, ΔsnR35 | / | / |
| Figure 4F | ΔLtv1, Gal:S3 | Ltv1_S/D + WT S3 | TEF413 for Ltv1_S/D and TEF 416 for S3 plasmids |
|  |  | Ltv1_S/D + S3_KK |  |
|  | ΔLtv1, ΔsnR35, Gal:S3 | Ltv1_S/D + WT S3 |  |
|  |  | Ltv1_S/D + S3_KK |  |
| Figure S3A | ΔLtv1, Gal:Tsr1, Gal:S20 | Ltv1_S/D + WT S20+ WT Tsr1 | TEF413 for Ltv1_S/D, TEF 415 for S20 plasmids and TEF 416 for Tsr1 plasmids. |
|  |  | Ltv1_S/D + S20_DE + WT Tsr1 |  |
|  |  | Ltv1_S/D + S20_EYER + WT Tsr1 |  |
|  |  | Ltv1_S/D + WT S20+ Tsr1_RK |  |
|  |  | Ltv1_S/D + S20_DE + Tsr1_RK |  |
|  |  | Ltv1_S/D + S20_EYER + Tsr1_RK |  |
|  | ΔLtv1, Gal:Rio2, Gal:S20 | Ltv1_S/D + WT S20+ WT Rio2 | TEF416 for Ltv1_S/D, TEF 413 for Rio2 plasmids and TEF 415 for S20 plasmids |
|  |  | Ltv1_S/D + S20_DE + WT Rio2 |  |
|  |  | Ltv1_S/D + S20_EYER + WT Rio2 |  |
|  |  | Ltv1_S/D + WT S20+ Rio2_loop |  |
|  |  | Ltv1_S/D + S20_DE + Rio2_loop |  |
|  |  | Ltv1_S/D + S20_EYER + Rio2_loop |  |
| Figure S3B | Gal:Tsr1 | WT Tsr1 | TEF416 |
|  |  | Tsr1_RK |  |
|  | ΔLtv1, Gal:Tsr1 | WT Tsr1 |  |
|  |  | Tsr1_RK |  |
|  | Gal:Rio2 | WT Rio2 | Cyc1 415 |
|  |  | Rio2_K105E |  |
|  |  | Rio2_loop |  |
|  | ΔLtv1, Gal:Rio2 | WT Rio2 |  |
|  |  | Rio2_K105E |  |
|  |  | Rio2_loop |  |
| Figure S3C | Gal:Tsr1 | WT Tsr1 | TEF416 |
|  |  | Tsr1_RK |  |
|  | Gal:Rio2 | WT Rio2 | Cyc1 415 |
|  |  | Rio2_K105E |  |
|  |  | Rio2_loop |  |
| Figure S3D | Gal:Rio2 | WT Rio2 | Cyc1 415 |
|  |  | Rio2_K105E |  |
|  |  | Rio2_loop |  |
|  |  | Rio2_Δloop |  |
| Figure S4A | BY4741 | / | / |
|  | ΔsnR35 | / | / |
| Figure S4C | Gal:Rio2 | WT Rio2 | Cyc1 415 |
|  |  | Rio2_K105E |  |
|  |  | Rio2_loop |  |
|  |  | Rio2_Δloop |  |
|  | ΔsnR35, Gal:Rio2 | WT Rio2 |  |
|  |  | Rio2_K105E |  |
|  |  | Rio2_loop |  |
|  |  | Rio2_Δloop |  |
|  | Gal:Tsr1 | WT Tsr1 | TEF416 |
|  |  | Tsr1_RK |  |
|  | ΔsnR35, Gal:Tsr1 | WT Tsr1 | TEF415 |
|  |  | Tsr1_RK |  |
| Figure S4D | BY4741 | / | / |
|  | ΔLtv1 | / | / |
|  | ΔsnR57 | / | / |
|  | ΔLtv1, ΔsnR57 | / | / |
